# The Medical Genome Reference Bank: Whole genomes and phenotype of 2,570 healthy elderly

**DOI:** 10.1101/473348

**Authors:** Mark Pinese, Paul Lacaze, Emma M. Rath, Andrew Stone, Marie-Jo Brion, Adam Ameur, Sini Nagpal, Clare Puttick, Shane Husson, Dmitry Degrave, Tina Navin Cristina, Vivian F. Silva Kahl, Aaron L. Statham, Robyn L. Woods, John J. McNeil, Moeen Riaz, Margo Barr, Mark R. Nelson, Christopher M. Reid, Anne M. Murray, Raj C. Shah, Rory Wolfe, Joshua R. Atkins, Chantel Fitzsimmons, Heath M. Cairns, Melissa J. Green, Vaughan J. Carr, Mark J. Cowley, Hilda A. Pickett, Paul A. James, Joseph E. Powell, Warren Kaplan, Greg Gibson, Ulf Gyllensten, Murray J. Cairns, Martin McNamara, Marcel E. Dinger, David M. Thomas

## Abstract

Population health research is increasingly focused on the genetic determinants of healthy ageing, but there is no public resource of whole genome sequences and phenotype data from healthy elderly individuals. Here we describe the Medical Genome Reference Bank (MGRB), comprising whole genome sequence and phenotype of 2,570 elderly Australians depleted for cancer, cardiovascular disease, and dementia. We analysed the MGRB for single-nucleotide, indel and structural variation in the nuclear and mitochondrial genomes. Individuals in the MGRB had fewer disease-associated common and rare germline variants, relative to both cancer cases and the gnomAD and UK BioBank cohorts, consistent with risk depletion. Pervasive age-related somatic changes were correlated with grip strength in men, suggesting blood-derived whole genomes may also provide a biologic measure of age-related functional deterioration. The MGRB provides a broadly applicable reference cohort for clinical genetics and genomic association studies, and for understanding the genetics of healthy ageing. This research has been conducted using the UK Biobank Resource under Application Number 17984.

## Introduction

Most developed nations face crises in health care associated with population ageing (Prince et al., 2015). Healthy ageing is a complex phenotype, influenced by both environmental and genetic factors (Lowsky et al., 2014). Healthy ageing – the absence of clinically significant, non-communicable disease or morbidity in old age – is distinct from longevity, which disregards quality of life. Healthy ageing captures the critical distinction between a long life with minimal impairment, and one bearing significant, costly, and potentially prolonged morbidity (Brooks-Wilson, 2013).

Relatively little is known about the genetic determinants of ageing that account for the broad spectrum of health states observed in older people. Genetic variation contributes to healthy ageing through pleiotropic effects on many diseases, immune responses, anthropomorphic, and behavioural phenotypes (Tobacco and Genetics Consortium, 2010; Wray et al., 2018). For example, alleles associated with behavioural phenotypes that contribute to a healthy lifestyle, such as avoidance of smoking, or propensity for regular exercise, might be anticipated to have an effect on healthy ageing. However, to date, Genome Wide Association Studies (GWAS) of common variants have only consistently associated the *APOE* and *FOXO3A* loci with lifespan and longevity (Brooks-Wilson, 2013), with *TERT* also implicated in ageing rate (Lu et al., 2018). Rare pathogenic variants that hasten the onset of common diseases associated with age, such as cancer, cardiovascular disease, and neurodegenerative disorders, might also be expected to be depleted in healthy aged individuals. While the single study using whole genome sequencing (WGS) in 511 healthy aged individuals confirmed the link with the *APOE* locus, and suggested depletion of polygenic risk for Alzheimer’s disease and coronary artery disease (Erikson et al., 2016), no evidence was found for depletion of rare pathogenic variation. Limited by sample size, these studies have focussed on single nucleotide variants and indels, while large scale structural variation remains unexplored. In addition, somatic variation such as clonal haematopoiesis (CH) is known to correlate with both age and susceptibility to disease (Genovese et al., 2014; Jaiswal et al., 2014). A synthesis of all forms of somatic and germline genomic variation is needed to inform our understanding of healthy ageing and disease susceptibility.

The advent of WGS is driving intense interest in mapping the genetic basis of disease, less than 50% of which is currently understood (Visscher et al., 2017). The missing heritability arguably resides in the total burden of both common and rare variation, structural variation untagged by simple polymorphisms, and their interactions (Zuk et al., 2012, 2014). WGS enables more comprehensive characterization of common, rare, and complex variation in human cohorts. The next few years will see the release of large-scale WGS studies in rare diseases and cancer, such as the 100,000 Genomes Project, and population studies like the UK Biobank. Maximising the analytic power of whole genome association studies using these cohorts will require well-phenotyped and high-quality control data. The concept of extreme phenotype sampling maximizes statistical power by comparing the extremes of phenotypes of interest (Barnett et al., 2013; Li et al., 2011; Zhou et al., 2016). We postulate that an elderly cohort depleted of the major common diseases constitutes a powerful and broadly applicable tool for genome-wide association studies of disease.

With this background, we undertook WGS of 2,570 elderly individuals with no reported history of cancer, cardiovascular disease, or neurodegenerative diseases up to age 70, to create the Medical Genome Reference Bank (MGRB) (Lacaze et al., 2018). For comparison, we have also undertaken whole genome sequencing of 344 young subjects, and 273 elderly individuals with cancer. These cohorts have been subjected to a broad spectrum, systematic analysis of germline and somatic variation within the nuclear and mitochondrial genomes, which we have linked to both chronologic age as well as frailty measures.

## Results

### Cohort characteristics and sequencing

We identified 2,926 individuals from the ASPREE study (McNeil et al., 2017), and Sax Institute’s 45 and Up Study (45 and Up Study Collaborators et al., 2008), who lived to at least 70 years of age without any history of cancer, cardiovascular disease, or dementia, confirmed either at baseline entry or study follow-ups (Lacaze et al., 2018). We sequenced all samples by WGS, mapping to build 37 of the human reference genome, and calling variants following GATK best practices. After exclusion of 356 samples that failed quality control and relatedness checks, 2,570 samples remained, forming the MGRB cohort (Table 1).

**Table 1:** Summary metrics for the MGRB well elderly cohort, sourced from the ASPREE or 45 and Up studies. Aggregate statistics are medians, with ranges in parentheses. Genetic background (ancestry) was determined from genotype data. Although blood was occasionally drawn at younger than 70 years, all individuals lived to at least 70 years without known cancer, cardiovascular disease, or dementia.

| *Measure* | *ASPREE* | *45 and Up* |
| --- | --- | --- |
| Individuals<br>(percent female) | 1,853<br>(48.2%) | 717<br>(59.3%) |
| Age at blood draw (years) | 79<br>(75 – 95) | 70<br>(64 – 91) |
| Height (m) | 1.65<br>(1.33 – 1.91) | 1.66<br>(1.37 – 1.91) |
| Mass (kg) | 74.5<br>(33.4 – 127.1) | 72.0<br>(36.0 – 147.0) |
| Mean sequencing depth<br>(genome-wide) | 38.0<br>(26.8 – 46.0) | 39.0<br>(27.3 – 45.5) |
| Genetic background |  |  |
| Non-Finnish European | 1,805 | 695 |
| South and Central American | 23 | 5 |
| South Asian | 14 | 6 |
| Finnish European | 10 | 7 |
| East Asian | 1 | 4 |

A broad diversity of genetic variation was found in the MGRB cohort. We identified 69,996,670 small variant loci in canonical chromosome contigs, with a call rate of 99.5%. Our small variant detection sensitivity was 99.3% and false positive rate 4.84 / Mbp, as assessed by comparing an internal RM 8398 sample against a gold standard (Zook et al., 2014). MGRB participants were primarily of non-Finnish European genetic background (Table 1, Supplementary Figure 1). Consistent with previous studies (Lek et al., 2016; Telenti et al., 2016), 51.8% of small variants were singletons, and 4.6% of loci were multi-allelic.

In addition to small scale variants, an average of 4,036 structural variants (SVs) per individual were observed, most commonly deletions (Supplementary Table 1, Supplementary Figure 2). In contrast to small variants, only 17% of structural variants were unique (Supplementary Table 2). Each individual carried an average of 4,264 mobile element insertions (MEI), predominantly of the ALU and L1 classes, consistent with a previous report (Chaisson et al., 2015), and most MEIs were copy number polymorphisms at known loci. However, on average 1,535 MEI events per individual were in regions of the reference genome not currently described as containing mobile elements. In summary, while small variants comprise the majority of genetic diversity in the MGRB, structural and mobile elements constitute a rich and understudied source of potentially disease-related variation.

### The well elderly carry clinically reportable genetic variation

Population genomic studies are contributing to the substantial revision of clinical interpretation of genetic variation thought to drive disease in some cases (Walsh et al., 2017). It is therefore clinically important to understand the frequency of variants currently considered pathogenic in a clinical context, but which are observed in well elderly individuals (Lacaze et al., 2017). To this end, we identified pathogenic variants that are considered clinically reportable as incidental findings under current American College of Medical Genetics (ACMG) guidelines (Kalia et al., 2016; Richards et al., 2015). Forty pathogenic or likely pathogenic heterozygous small variants were identified, with 28/2,570 (1.1%) individuals carrying dominantly acting variants linked to disease (Table 2, Supplementary Data File 1). We sought further evidence of disease phenotypes in individuals carrying relevant pathogenic variants from the ASPREE cohort. We did not identify personal histories of breast or colorectal cancer in individuals harbouring *BRCA2*, *MSH2,* or *PMS2* mutations; cardiac arrest or strokes in individuals harbouring *DSG2*, *DSP*, *KCNH2*, *KCNQ1*, *MYBPC3*, *MYL3*, and *SCN5A* mutations; or elevated blood lipid levels in *APOB* carriers.

**Table 2:** Counts of clinically significant small variation in the MGRB for all genes in the ACMG SF 2.0 set. Abbreviations: ARVC, Arrhythmogenic right ventricular cardiomyopathy; CPVT, Catecholaminergic polymorphic ventricular tachycardia; HCM, Hypertrophic cardiomyopathy; DCM, Dilated cardiomyopathy; VA, Ventricular arrhythmia.

| *Condition* | *Gene* | *Carriers* |
| --- | --- | --- |
| Cancer | *BRCA2* | 4 (2 female) |
|  | *MSH2* | 1 |
|  | *MSH6* | 1 |
|  | *PMS2* | 3 |
| Neurofibromatosis | *NF2* | 1 |
| ARVC | *DSG2* | 1 |
|  | *DSP* | 3 |
| CPVT | *RYR2* | 1 |
| HCM, DCM | *MYBPC3* | 2 |
|  | *MYL3* | 1 |
|  | *TNNI3* | 1 |
| Hypercholesterolemia | *APOB* | 5 |
| Long QT, VA | *KCNH2* | 1 |
|  | *SCN5A* | 1 |
| Marfan syndrome | *MYH11* | 1 |
| Malignant hyperthermia | *RYR1* | 1 |
| TOTAL |  | 28 |

Cancer-associated genotypes are dependent on stochastic factors which may account for variable penetrance, while anaesthetic-associated malignant hyperthermia linked to loss of function variation in *RYR1* is contingent on environmental exposure. We specifically sought, but did not find, evidence of cardiovascular disease history or related clinical phenotypes in carriers of variants linked to atrial fibrillation, cardiomyopathy and hypertension. Notably, no genotypes predicted to cause severe childhood-onset diseases were identified (Chen et al., 2016); the single *RYR2* variant detected was a truncation not expected to cause autosomal dominant catecholaminergic polymorphic ventricular tachycardia. In five individuals, variants were noted in *PCSK9* that are predicted to be protective against high blood cholesterol (Langsted et al., 2016), comparable to the rate observed in the gnomAD non-Finnish European cohort (Fisher exact test, p = 0.37). Four SVs were found that may disrupt the coding sequence of genes associated with cancer and cardiovascular health (Supplementary Table 3), comprising 10% of potentially pathogenic variation in genes considered reportable by the ACMG.

### Rare and common risk variants are depleted in the well elderly

One of the primary purposes of the MGRB is to serve as a genetic risk-depleted control cohort for studies of the common causes of morbidity and mortality. To test its utility, we compared the rates of pathogenic variants in tumour suppressor genes between the 717 MGRB individuals from the 45 and Up Study, with 269 demographically-matched cancer cases from the same study (45 and Up Study; Supplementary Table 4). Considering all cancers in aggregate, the MGRB samples were significantly depleted for pathogenic alleles in tumour suppressor genes relative to cancer cases, with 2 of 717 controls carrying pathogenic tumour suppressor variants compared to 12 of 269 cancer cases (Figure 1a, odds ratio 0.060, 95% confidence interval 0.0065 to 0.27, p < 0.001, Fisher’s exact test). In addition to all cancers, we specifically examined colorectal cancer due to its high incidence in our case set, and well-defined genetic risk (De Rosa et al., 2015). The MGRB samples were significantly depleted for rare pathogenic variation in the *APC*, *MLH1*, *MSH2*, *MSH6*, *PMS2*, and *SMAD4* genes, relative to colorectal cancer cases (Figure 1a, 1 of 717 MGRB with pathogenic variants, versus 2 of 40 cancer cases, odds ratio 0.027, 95% CI 0.001 to 0.53, p = 0.008). We did not detect a difference in the rate of rare coding loss of function variants in the MGRB relative to the gnomAD non-Finnish European cohort, either genome-wide, or in genes associated with cardiovascular disease or cancer, potentially due to technical factors dominating differences in rare variant patterns between cohorts (8 tests, all p > 0.08).

**Figure 1:**
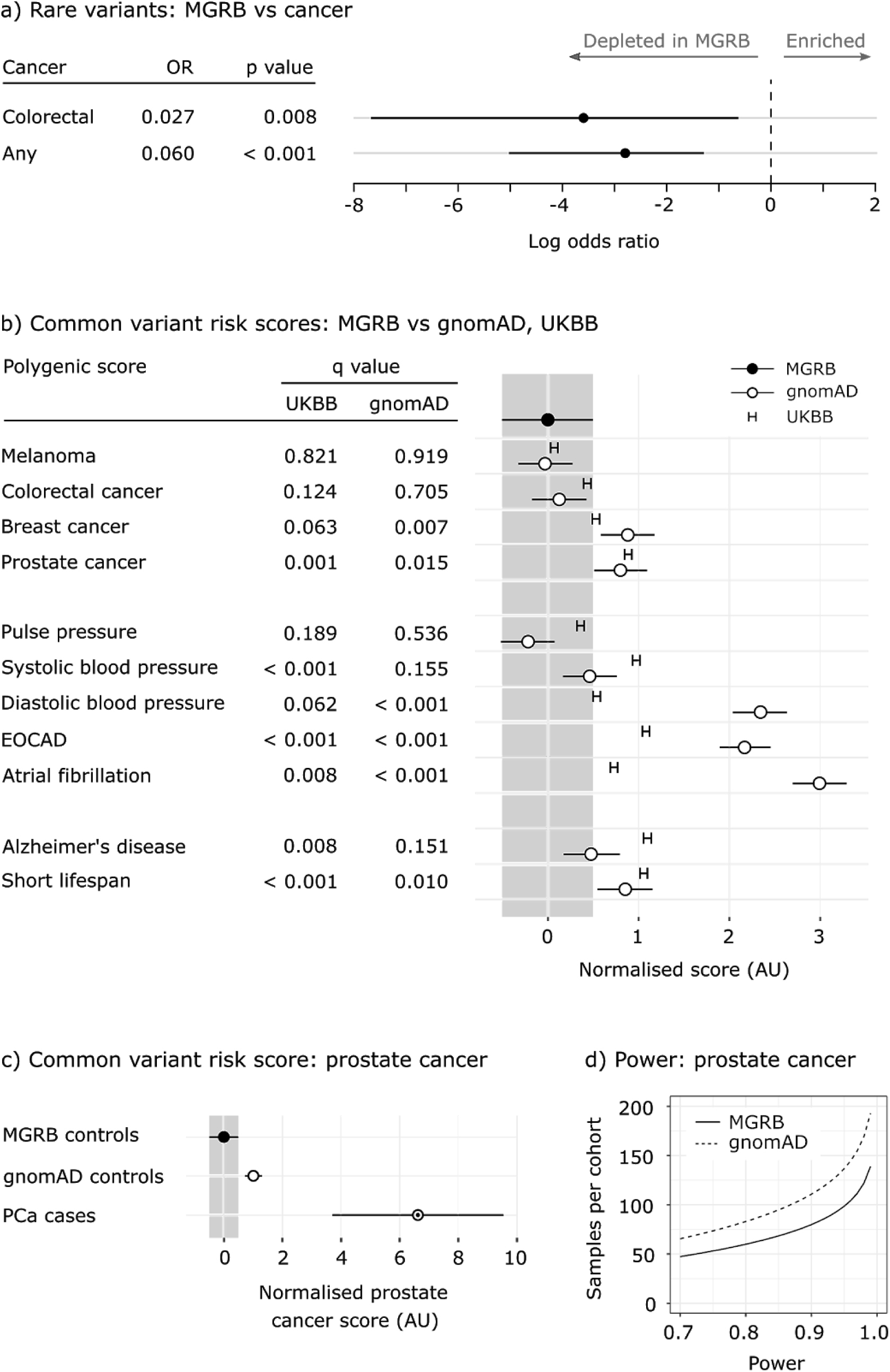
The MGRB is depleted for genomic risk relative to reference and disease cohorts. a) the rate of rare pathogenic variants in tumour suppressor genes is lower in MGRB than in a cohort of cancer cases (log odds for an individual to carry a pathogenic tumour suppressor variant shown). b) the MGRB also has lower polygenic score (PS) estimates for a range of phenotypes, when compared to the gnomAD non-Finnish European population and the UK BioBank samples. MGRB is the reference in (b), with PS mean set at zero; bootstrap 95% confidence intervals are shown for each PS. q-values represent false discovery rate estimates by the Benjamini-Hochberg method (Benjamini and Hochberg, 1995). c) the MGRB has lower PS compared to prostate cancer cases, here considering only samples from the 45 and Up Study. d) for any given sample size, the MGRB has greater statistical power to detect PS difference from a case cohort than to gnomAD, demonstrated here for prostate cancer. AU: arbitrary units.

We next sought evidence for depletion of common disease-associated variation in the MGRB, relative to the gnomAD and UKBB datasets. Although SNP allele frequencies were highly concordant across all three cohorts (Supplementary Figure 3), the MGRB cohort was significantly depleted for alleles specifically associated with risk of cancer, cardiovascular disease, and neurodegenerative disease (Supplementary File 2, 698 loci, odds ratios 0.38 vs gnomAD, 0.47 vs UKBB, both p < 1.1×10^−6^, Fisher’s exact test). This enrichment of protective alleles was specific to the clinical phenotypes excluded from MGRB, and was not observed for negative control loci linked to anthropometric (449 loci, both p > 0.69) or behavioural (575 loci, both p > 0.55) traits.

The aggregate burden of common disease-related variants within individuals can be summarised in a polygenic score (PS). We constructed polygenic predictors for a range of phenotypes measured or depleted in the MGRB, and compared PS distributions between MGRB, the gnomAD non-Finnish European reference cohort, the UK BioBank (UKBB), and the 45 and Up Study cancer cohort. Significant depletion in PS was observed in MGRB for eight of the eleven scores tested (Figure 2b). Notably, a PS associated with short lifespan (Deelen et al., 2014) was significantly depleted in MGRB relative to gnomAD and UKBB, consistent with the MGRB healthy elderly phenotype. MGRB individuals were significantly depleted for prostate cancer risk relative to both gnomAD and prostate cancer cases, indicating that MGRB is an extreme depletion cohort for prostate cancer polygenic risk (Figure 2c). Critically, for the extreme phenotype sampling hypothesis, the use of the MGRB as a control cohort reduced the sample size required to reach a given target power by approximately 25% by comparison with the widely-used gnomAD dataset (Figure 1d).

**Figure 2:**
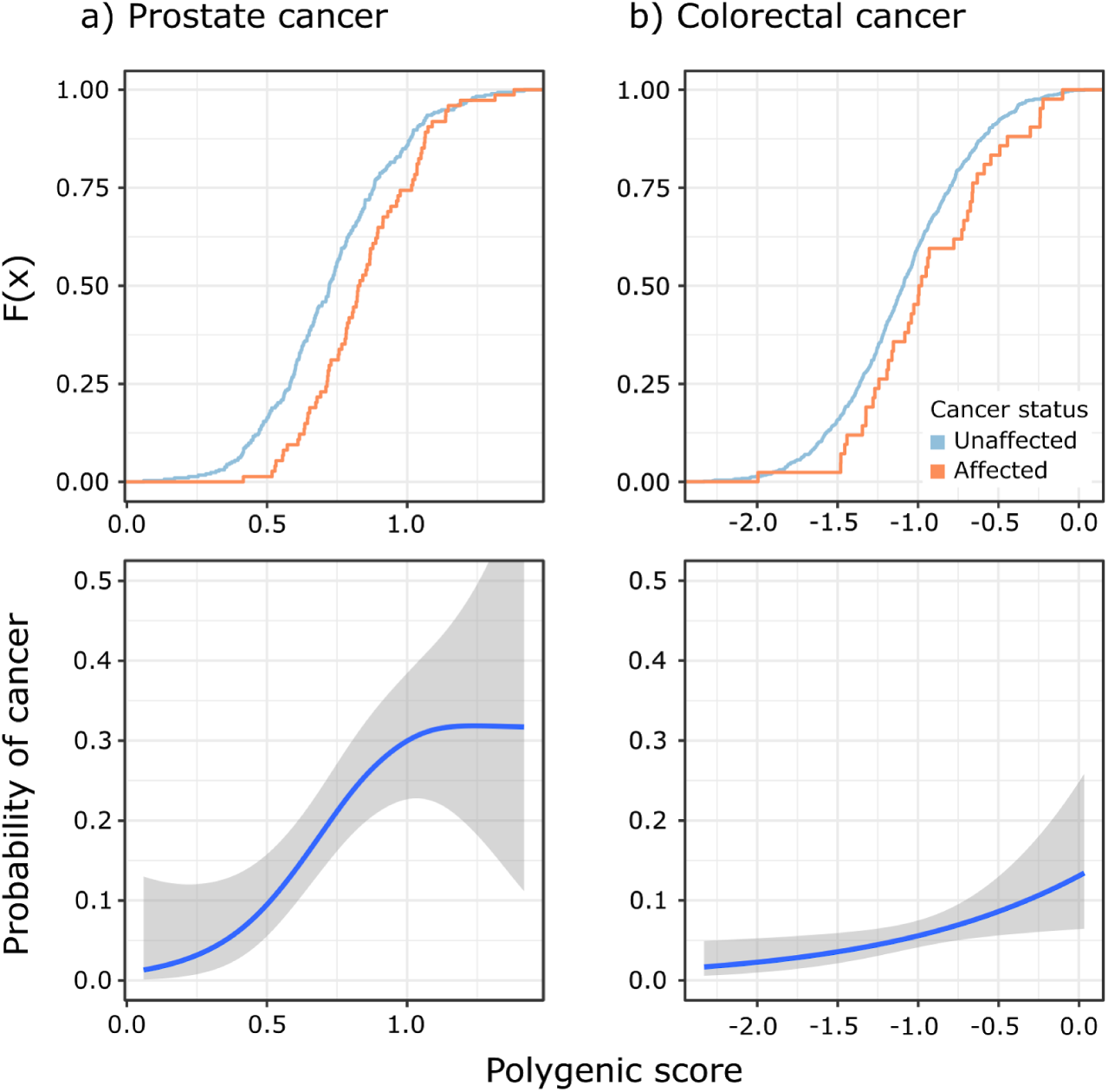
Polygenic risk is strongly related to cancer diagnosis risk. Cumulative distribution functions (top panels) and associated probability of cancer diagnosis by age 70 (bottom panels) are shown for both prostate cancer (a) and colorectal cancer (b). Unaffected individuals are MGRB men (prostate), or all MGRB individuals (colorectal) and were completely cancer-free up to age 70; affected individuals were sourced from the 45 and Up Study cancer cohort and had recorded evidence of the relevant cancer diagnosis prior to age 70. Polygenic scores were computed based on reported loci and model coefficients (Hoffmann et al., 2015; Schumacher et al., 2015). Fits are from logistic regression using a GCV-penalised thin plate spline smooth; bands denote 95% confidence intervals around the mean.

In addition to the allele frequency-based comparisons above, the availability of individual genotypes for the MGRB and 45 and Up Study cancer cohorts enabled the direct evaluation of the influence of PS on cancer risk. We first confirmed that our polygenic scoring method estimated individual height using published loci (Wood et al., 2014): height PS was significantly predictive of measured height, with a slope of 4.5 cm per polygenic score unit, complete model R^2^ = 0.62, polygenic score partial R^2^ = 0.14, n = 2,537 (Supplementary Figure 5). We then compared the distribution of PS for prostate, colorectal, and melanoma skin cancer between the 45 and Up cancer-free cases in the MGRB, and individuals from the 45 and Up cohort with these cancers (Supplementary Table 4). Consistent with the relative depletion of rare cancer variants in the MGRB observed above, MGRB individuals had significantly lower polygenic risk scores than cases for prostate cancer (p < 0.001, Figure 2a) and colorectal cancer (p = 0.022, Figure 2b), but not melanoma. The contribution of PS to cancer-specific risk was significant: by age 70, individuals with a cancer PS in the top 5% of MGRB had a 7.7-fold increased odds for prostate cancer, and a 3.6-fold increased odds for colorectal cancer, relative to individuals with a score in the bottom 5%.

### Clonal somatic variation is detectable by WGS

In addition to its use as a surrogate for the germline, peripheral blood DNA carries somatic variation reflecting the life history and health state of the donor. Clonal haematopoiesis of Indeterminate Potential (CHIP) occurs in at least 10% of individuals over the age of 65 years, evidenced by somatically acquired SNVs (Genovese et al., 2014; Young et al., 2016). Previous studies have used deep whole-exome or targeted sequencing to identify CHIP, which lacks sensitivity to detect the SVs commonly observed in myelodysplasia and leukemia. Low depth WGS, a powerful tool for measuring SV, has not been applied to the detection of CHIP. Here we estimate the burden of cancer-associated somatic variation in peripheral blood DNA using whole genome data in the MGRB cohort.

In total, 184/2,570 (7.2%) of MGRB individuals displayed evidence of CHIP, with SNVs associated with overgrowth and neoplasia observed in more than 10% of reads. Predominantly nonsense mutations (96%), these variants were most commonly seen in *TET2* (47 individuals), *DNMT3A* (23), or *ASXL1* (11). We also observed known gain-of-function missense variants in *JAK2* V617F (9 individuals), *NRAS* G12D (1), a dominant negative allele in *DNMT3A,* R882H (1) (Russler-Germain et al., 2014), and a putative loss-of-function variant in *TP53*, C275Y (1). *JAK2* V617F is a recognised driver of myeloproliferative disorders (Percy and McMullin, 2005), which are also associated with *ASXL1* loss (Gelsi-Boyer et al., 2012), and *TET2* and *DNMT3A* loss-of-function variants are frequent in CHIP (Buscarlet et al., 2017). In total, the blood of 91/2,570 (3.5%) cancer-free MGRB individuals carried deleterious small variation in at least one of these four genes, and 13 individuals had multiple deleterious mutations in this gene set. We next sought evidence for subclonal copy number variation (CNV). 1975 of 2570 MGRB individuals were successfully fit to a subclonal CNV model; of these 55 (2.8%) showed evidence of subclonal CNV, as determined by the presence of an aneuploid lineage representing more than 10% of nucleated blood cells (Supplementary Figure 6). In total, 9.2% (95% CI 7.9 to 10.5%) of MGRB samples demonstrated evidence of CHIP by either SNV or CNV, consistent with results from deep WES (Jaiswal et al., 2014). In sum, subclonal blood DNA changes are detectable from WGS at routine read depths used for germline purposes, providing a quantitative fingerprint of age-related somatic events.

### Age-related mitochondrial load is associated with grip strength

As well as CHIP, ageing is associated with telomere shortening (von Zglinicki and Martin-Ruiz, 2005), somatic Y chromosome loss, decreased mitochondrial copy number, and increased mitochondrial heteroplasmy (Kennedy et al., 2013; Wachsmuth et al., 2016). We therefore studied the relationship of age to telomere length, mitochondrial copy number and variation, Y copy number in males, a somatic mutation signature linked to ageing (Alexandrov et al., 2015), and CHIP. Using standard depth WGS data from multiple cohorts, consistent patterns of change with age were observed across all six somatic metrics (Figure 3a-f). Compared to a population of younger individuals (the ASRB cohort, median age 40, Supplementary Figure 4), the MGRB, despite being ascertained on the basis of healthy ageing, was still associated with shorter telomere lengths, increased somatic mutation burden, and decreased Y chromosome and mitochondrial copy number (Table 3). Interestingly, there were differences between each cohort in the relationship with age, with apparent stabilisation of telomere length in the elderly cohorts past approximately 70 years, compared to the expected progressive shortening with increasing age observed in the younger ASRB cohort. In addition, while mitochondrial copy number/nuclear genome was stable up to age 60, significant declines were observed in the older age groups. The rate of change was significantly different between the young (ASRB) and aged (MGRB) cohorts (5 likelihood ratio tests on linear fits, Holm correction, all p < 0.003), while the rate of change of the two aged cohorts was consistent across all measures (5 likelihood ratio tests, Holm correction, all p > 0.28). Taken together, these results are suggestive of altered kinetics of age-related somatic mutation in the elderly compared to younger populations, although we note longitudinal measurements will be necessary to definitively establish this relationship.

**Figure 3:**
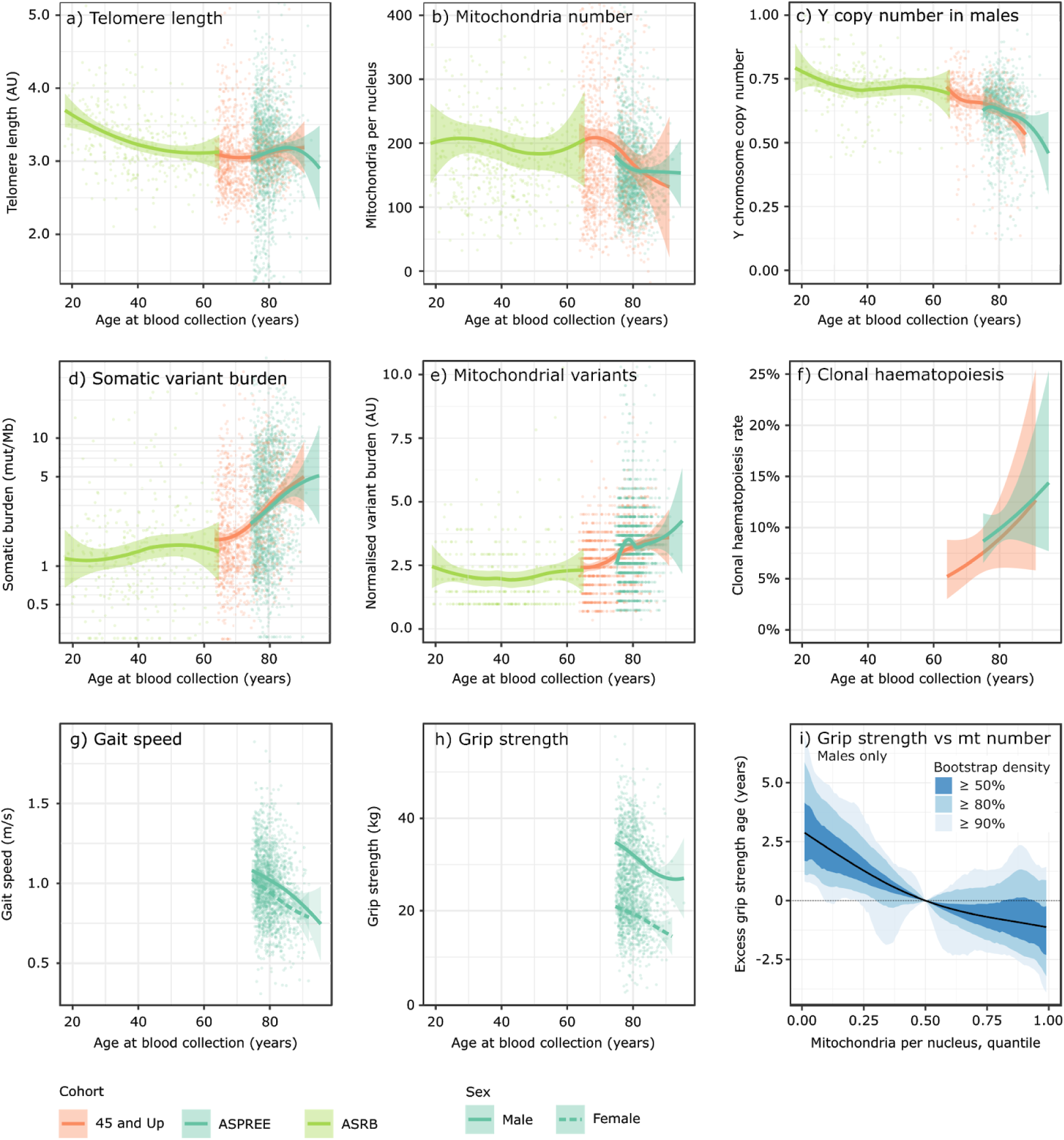
Age-related somatic changes detectable in blood DNA by whole genome sequencing are associated with two measures of physical function. Across multiple cohorts, a consistent decrease with age is observed for telomere length (a), mitochondria per nucleus (b), and Y copy number in males (c). In contrast, advanced age is associated with an increase in somatic mutation burden (d,e) and the fraction of samples with detectable clonal haematopoiesis (f), as well as a decrease in the key physical function measures gait speed (g) and grip strength (h). The count of mitochondria per nucleus is significantly related to grip strength beyond age alone in men, as indicated by the change in effective age as judged by grip strength with varying mitochondria count (i). For (a-c,g,h) individual measurements corrected for cohort batch effect are shown with LOESS smooths, and for (d) a logistic fit was used. Bands around estimates delimit 99% confidence intervals for the mean.

**Table 3:** The rates of somatic measure change with age are different between middle-aged and old individuals. Numbers show the rate of change of each somatic measure with age in the middle-aged ASRB cohort (median age 40), and the older MGRB cohorts (median age 70 or older). Changes are significantly different between the younger ASRB and older MGRB cohorts, and consistent within the two older MGRB cohorts. Linear model slopes as change per decade are reported for each of five somatic measures in each cohort, with 95% Wald confidence intervals. Values significantly different from zero are represented in bold. Note that somatic burden and mitochondrial count per nucleus are reported on the natural logarithm scale. N.D.: Not determined due to data use agreement constraints.

| *Measure* | *Cohort* |  |  |
| --- | --- | --- | --- |
|  | *ASRB* | *45 and Up Study* | *ASPREE* |
| Individuals | 344 | 717 | 1853 |
| Percent female | N.D. | 59.3% | 48.2% |
| Median age<br>(range) | 40<br>(18 - 65) | 70<br>(64 - 91) | 79<br>(75 - 95) |
| Telomere length<br>(AU/decade) | **-0.115**<br>[-0.157, -0.073] | 0.040<br>[-0.010, 0.090] | **0.115**<br>[0.035, 0.196] |
| Mitochondria count<br>(log<sub>10</sub> mt/nucleus/decade) | -0.004<br>[-0.018, 0.010] | **-0.046**<br>[-0.065, -0.027] | **-0.038**<br>[-0.059, -0.017] |
| Y copy number in males<br>(Y chromosomes/nucleus/decade) | -0.011<br>[-0.022, 0.001] | **-0.050**<br>[-0.068, -0.033] | **-0.043**<br>[-0.065, -0.021] |
| Somatic variant burden<br>(log<sub>10</sub> variants/Mb/decade) | 0.038<br>[-0.002, 0.079] | **0.207**<br>[0.167, 0.247] | **0.228**<br>[0.173, 0.282] |
| Mitochondrial variants<br>(mt variants/decade) | 0.051<br>[-0.177, 0.278] | **1.665**<br>[1.315, 2.015] | **0.893**<br>[0.195, 1.591] |

We next considered whether individual age-related genomic measures may reflect physical function status, independently of chronologic age. To address this question, somatic changes in MGRB samples from the ASPREE cohort were studied in the context of age, grip strength, and gait speed, all representing key predictors of age-related morbidity (Chainani et al., 2016; Dudzińska-Griszek et al., 2017; Syddall et al., 2003). As expected, grip strength and gait speed both consistently decreased with age in both genders (Figure 3g,h). To correct for the strong influence of age on all measures, a conditional analysis was performed to explore whether any somatic measures were associated with physical function even when age is taken into account. Intriguingly, we found that grip strength was positively correlated with the count of mitochondria per nuclear genome, but only in males (two-stage test, first stage p = 0.051, validation p = 0.036).

To illustrate the magnitude of effect of mtDNA copy number on grip strength in men, we modelled the change in “effective age” as determined by grip strength, as a function of mtDNA copy number. This revealed that men with an mtDNA copy number in the lowest 5% for their age have the same grip strength as men with average mtDNA levels, but who are 2.5 years older (Figure 3i).

## Discussion

Understanding the genetic underpinnings of healthy ageing is as important as, and relevant to, understanding the genetic basis of disease. The next decade will see the fruits of population-scale sequencing programs, much of which will be aimed at understanding the genetic origins of disease. To realise this mission, we need to understand the spectrum of genetic variability in the healthy, and whole genome data sets of healthy controls will be essential to identify genetic variation unrelated to disease (Manrai, Patel, Ioannidis, JAMA, 2018). To this end we created the MGRB, a whole-genome sequencing resource of deeply-phenotyped aged individuals (Lacaze et al., 2018).

Although depletion of some common disease-related alleles has been reported in the healthy aged (Brooks-Wilson, 2013; Deelen et al., 2014; Erikson et al., 2016), the MGRB reveals a striking depletion in disease-associated common and rare variation, relative to both affected cases, as well as datasets frequently used as controls in genetic studies, but not specifically depleted for disease phenotypes. In addition, the MGRB was enriched for protective alleles linked to healthy ageing. Our data also substantiate the premise that extreme phenotype enrichment can enhance statistical power in case:control genetic studies (Li et al., 2011) (Figure 1c,d).

Despite being healthy, over 1% of the MGRB still carry pathogenic small variants that are clinically-reportable under current ACMG guidelines (Table 2), consistent with previous observations (Erikson et al., 2016). A detailed review of individual phenotypes from a subset of mutation carriers excluded even subclinical manifestations of the expected disorders.

These data suggest that many apparently pathogenic small variants have variable penetrance, echoing a theme emerging from population genomic studies. Additionally, several rare structural variants were identified that may abrogate function of clinically-reportable genes (Supplementary Table 3). Future studies using whole genome-based data will benefit from the MGRB in quantitating the contribution of structural variation to ageing and disease. These observations suggest the MGRB may provide a filter for rare variants currently thought pathogenic in a clinical context.

The ageing process is accompanied by the emergence of somatic mutations in tissues other than blood, mitochondrial depletion and heteroplasmy, and progressive telomere shortening (Acuna-Hidalgo et al., 2017; Genovese et al., 2014; Jaiswal et al., 2014; Martincorena et al., 2015). We developed a suite of methods to detect these age-related changes using 30X whole-genome sequencing, and applied it to the elderly MGRB and a younger cohort.

Telomere shortening itself may directly increase the likelihood of neoplasia (Artandi et al., 2000), while oncogenic mutations in genes such as *TP53* may rescue the effects of telomere loss (Chin et al., 1999). Telomere dysfunction has been associated with impaired mitochondrial function (Sahin et al., 2011), linking these genomic features of ageing. Many of the somatic changes of ageing observed in the MGRB are associated with marrow stem cell depletion (de Haan and Van Zant, 1999), consistent with studies in telomerase-deficient mice (Lee et al., 1998).

Interestingly, we observed a shift in the age trajectory of multiple somatic metrics in the elderly compared to younger individuals, coincident with the emergence of clonal hematopoiesis. It is paradoxical that, in this and other cohorts, somatic clonal expansion driven by oncogenic mutations appears compatible with normal organ function (Genovese et al., 2014; Jaiswal et al., 2014; Martincorena et al., 2015). It is even possible that neoplastic events, such as telomere stabilisation, loss of tumor suppressor genes, or acquisition of oncogenic kinase mutations, might increase clonogenic efficiency of an ageing marrow stem cell compartment. Some support for this concept, reminiscent of antagonistic pleiotropy (Williams, 1957), comes from mice carrying a hypermorphic form of *Trp53,* in which protection from neoplasia was accompanied by accelerated hematopoietic ageing and diminished marrow reserve (Dumble et al., 2007; Tyner et al., 2002). If true, these findings suggest that strategies that suppress tumor formation may accelerate ageing.

We observed an intriguing link between somatic burden and decline in physical function, providing a potential measure of what distinguishes individuals sharing the same age, but different physical function status. The relative depletion of mitochondria per leukocyte appeared to be associated with reduced grip strength in males, after adjustment for age. This finding is consistent with evidence that mitochondrial dynamics are strongly involved in ageing and function, particularly in males (Latorre-Pellicer et al., 2016; Wachsmuth et al., 2016). We note that our power to detect such an effect is low when using a well elderly cohort, but believe there will be great interest in deriving quantitative measures of biological ageing from standard-depth whole genome sequencing.

Although the largest cohort of healthy elderly whole genomes amassed to date, the MGRB is still subject to limitations as a research and clinical tool. The investigation of extremely rare variants is limited by the MGRB’s size, and complicated by batch effects in rare variant calls (Tom et al., 2017). Furthermore, the MGRB comprises almost exclusively white Australians, and follow-up studies will be required to assess the spectrum of genetic variation in the healthy elderly from other backgrounds. The MGRB was recruited on the basis of a restricted definition of healthy ageing, being depletion of cancer, cardiovascular disease, and dementia, and MGRB participants do bear other morbidities. However we note that the deep phenotype which accompanies the MGRB enables more focussed participant selection and the construction of for-purpose subset cohorts, making the MGRB of value as a universal control that can be depleted of any measured phenotype. Finally, although we observed associations between somatic measures and age that are suggestive of changes in ageing kinetics, this cannot be definitively established using our cross-sectional study design. Further studies with longitudinal samples will be required to verify our hypothesis of altered ageing kinetics, for which the methodology established here will be valuable.

Quantitative biomarkers of age may provide a summative metric of diverse genetic and environmental effects on health. Interpreted as endophenotypes, such biomarkers show promise to increase our ability to detect genetic patterns associated with ageing rate (Lu et al., 2018), but their true utility may be greater still as clinical tools in their own right. By encoding the aggregate influence of complex and potentially unmeasurable genetic and environmental effects over the life of an individual, biomarkers of age may represent health and disease risk with greater fidelity than external indicators such as calendar age or functional state.

Particularly with respect to cancer, the DNA-based measures of biological age we have demonstrated here may represent an individual’s underlying mutation rate, and therefore true cancer risk, due to combined genetic and environmental factors. This biomarker-centric perspective on cancer risk represents a synthesis and simplification of the traditional genotype- and environment-centric views, and we believe is a promising lens through which to consider disease risk, and differentiate normal compensatory changes associated with ageing, from those that precede malignancy.

## Acknowledgements

Whole genome sequencing of the MGRB, ASRB, and 45 and Up cancer cohort was undertaken at the Kinghorn Centre for Clinical Genomics, which is supported by the Kinghorn Foundation. Sequencing of these cohorts was funded through the NSW Genomics Collaborative Grants scheme from the NSW Office for Health and Medical Research.

Processing and warehousing of genomic data employed resources and services from the National Computational Infrastructure (NCI), which is supported by the Australian Government.

The authors acknowledge the ASPREE Healthy Ageing Biobank, ASPREE Investigator Group and ASPREE Collaborating Practitioners listed on www.aspree.org. ASPREE was funded by the National Institute on Aging and the National Cancer Institute at the National Institutes of Health (grant number U01AG029824); the National Health and Medical Research Council of Australia (grant numbers 334047, 1127060); the Victorian Cancer Agency and Monash University (Australia). The ASPREE Healthy Ageing Biobank was supported by the Commonwealth Scientific and Industrial Research Organisation (Australia), the Victorian Cancer Agency (Australia) and Monash University (Australia). We acknowledge the dedicated and skilled staff in Australia and the U.S. for the conduct of the ASPREE trial, and the ASPREE participants who willingly volunteered.

This research was completed using data collected through the 45 and Up Study (www.saxinstitute.org.au). The 45 and Up Study is managed by the Sax Institute in collaboration with major partner Cancer Council NSW; and partners: the National Heart Foundation of Australia (NSW Division); NSW Ministry of Health; NSW Government Family & Community Services – Ageing, Carers and the Disability Council NSW; and the Australian Red Cross Blood Service. We thank the many thousands of people participating in the 45 and Up Study.

Genomic analysis of the Australian Schizophrenia Research Bank (ASRB) was supported by a New South Wales Health, Collaborative Genomics grant program (MC, MG, VC), a NARSAD Independent Investigator Grant (MC) and National Health and Medical Research Council (NHMRC) project grants (1067137, 1147644, 1051672). Samples were also collected by the ASRB with the support of the NHMRC, the Pratt Foundation, Ramsay Health Care, and the Viertel Charitable Foundation. The ASRB was supported by the Schizophrenia Research Institute (Australia), utilizing infrastructure funding from NSW Health and the Macquarie Group Foundation.

This research has been conducted using the UK Biobank Resource under Application Number 17984.

D.M.T is the recipient of an NHMRC Principal Research Fellowship (RegKey:1104364). M.J.Cairns was supported by an NHMRC Senior Research Fellowship (#1121474). M.J.Cowley was supported by a NSW Health Early-Mid Career Fellowship. J.R.A. was supported by University of Newcastle RHD and an Emlyn and Jennie Thomas Postgraduate Medical Research Scholarship. M.E.D., C.P., W.K., D.D., and S.H. were supported by the Kinghorn Foundation.

The authors thank Prof. Christopher Goodnow, Prof. Diane Fatkin, Dr Catherine Vacher for helpful comments and discussion, Dr Eleni Giannoulatou for the atrial fibrillation polygenic score coefficients, and Verity Hodgkinson for biospecimen management.

## Author Contributions

Conceptualization, M.P., M.E.D., and D.M.T.;

Methodology, M.P., E.M.R., J.E.P., C.P., and D.M.T.;

Software, M.P., E.M.R., and A.L.S;

Formal Analysis, M.P., E.M.R., A.L.S., M.R., C.P., M.J.Cowley, and P.A.J.;

Investigation, V.F.S K, and H.A.P.;

Resources, P.L., A.A., M.B., M. McN., E.B., M.R.N., C.M.R., A.M.M., R.C.S., R.W., M.J.Cairns., H.M.C, J.R.A, C.F., M.J.G., V.J.C., S.N., G.G., R.L.W., U.G., S.R., and J.J.McN.;

Data Curation, M.P., P.L., M.B., M. McN., and T.J.N C;

Writing -- Original Draft, M.P.;

Writing -- Review &

Editing, M.P., P.L., and D.M.T.;

Visualization, S.H., D.D., and W.K.;

Project Administration, A.S., and M-J.B.;

Funding Acquisition, M.E.D., and D.M.T.;

Supervision, M.E.D., and D.M.T.

## STAR Methods

### Experimental Model and Subject Details

#### Human subjects

Participants of the MGRB were consented through the biobank programs of the ASPREE and 45 and Up studies following protocols described previously (45 and Up Study Collaborators et al., 2008; Lacaze et al., 2018). At the time of blood collection, each participant was aged 60 years or older.

Samples from the ASPREE study were from individuals aged 75 years or older at time of enrolment, with no reported history of any cancer type, no clinical diagnosis of atrial fibrillation, no serious illness likely to cause death within the next 5 years (as assessed by general practitioner), no current or recurrent condition with a high risk of major bleeding, no anaemia (haemoglobin > 12 g/dl males, > 11 g/dl females), no current continuous use of other antiplatelet drug or anticoagulant, no systolic blood pressure ≥180 mm Hg and/or a diastolic blood pressure ≥105 mm Hg, no history of dementia or a Modified Mini-Mental State Examination (3MS) score ≤77 (Teng and Chui, 1987), and no severe difficulty or an inability to perform any one of the 6 Katz basic activities of daily living (Katz and Akpom, 1976).

Samples from the 45 and Up Study were from individuals with no self-reported history of cancer, heart disease, or stroke. Neurological disease was not explicitly excluded, but participants were required to correctly self-complete a health survey at enrolment. We confirmed no record of cancer diagnosis in the NSW Central Cancer Registry, and no record of cancer diagnosis in the NSW Admitted Patient Data Collection, for all 45 and Up Study individuals in the MGRB.

Participants in the Australian Schizophrenia Research Bank (ASRB) were recruited through a national media campaign and consented to data and sample collection genomic analyses as previously described (Loughland et al., 2010).

#### Ethics

The ASPREE Biobank study was approved by the Monash University Human Research Ethics Committee, and subsequent whole genome sequencing of Australian ASPREE participants was approved by the Alfred Hospital Ethics Committee. The use of 45 and Up Study samples in the MGRB is covered by ethics approvals from the University of New South Wales Human Research Ethics Committee and the NSW Population & Health Services Research Ethics Committee. The use of the ASRB data was approved by the University of Newcastle Human Ethics Research Committee.

### Method Details

#### Sample collection and processing

For ASPREE participants of the MGRB, peripheral blood samples were processed to buffy coat within 4 hours of collection, then stored at −80°C. DNA was later purified from buffy coat via magnetic bead extraction (Qiagen).

For 45 and Up Study participants of the MGRB, peripheral blood samples were refrigerated at 4°C and processed to buffy coat within 48 hours of collection. Buffy coat was stored at −80°C, and DNA purified via column extraction (Qiagen).

ASRB participant PBMCs were extracted from whole blood by centrifugation in Lymphoprep (Vital Diagnostics). Genomic DNA (gDNA) was extracted from PBMCs using salt extraction and quantified by PicoGreen assay (Life Technologies). The integrity of gDNA was determined by agarose gel electrophoresis prior to sequencing.

#### Sequencing

Whole genome sequencing of the MGRB, 45 and Up cancer, and ASRB cohorts was performed on Illumina HiSeq X sequencers at the Kinghorn Centre for Clinical Genomics (KCCG), Sydney, using paired-end Illumina TruSeq Nano DNA HT libraries and v2.5 clustering and sequencing reagents. 100 / 2,926 MGRB and 85 / 520 ASRB samples were sequenced to high depth (3 HiSeq X lanes per sample, equivalent to approximately 105X human genome), the remainder were sequenced to one lane per sample. Only one lane of data per sample was used to create the MGRB Phase 2 data release.

### Quantification and Statistical Analysis

#### Sequence alignment and processing

All sequence data generated at the KCCG were processed following the Genome Analysis Toolkit (GATK) best practices (Van der Auwera et al., 2013). We first defined a custom reference genome tailored to Illumina HiSeq X sequencers, being the 1000 Genomes Phase 3 decoyed version of build 37 of the human genome (The 1000 Genomes Project Consortium, 2015), with an added contig of NC_001422.1 to act as a decoy for the HiSeq-specific ΦX174 sequence spike-in. Reads were aligned to this reference using bwa 0.7.15 mem in paired mode, and duplicates marked with biobambam2 2.0.65 bamsormadup, with a minimum optical pixel distance of 2,500. All other parameters for both bwa and bamsormadup were left at defaults. For high-depth samples run on multiple sequencing lanes, data merging was performed at this point using samtools 1.5. Indel realignment and base quality score recalibration of mapped reads were performed using GATK 3.7-0 and best practices parameters; unmapped reads were left unmodified. GATK HaplotypeCaller was used to generate g.vcfs from all single-lane realigned and recalibrated BAMs using recommended parameters. Pipeline steps were accelerated using GNU parallel 20170722 (Tange, 2011).

#### Locus confidence tiers

We defined locus confidence tiers for WGS genotyping on the basis of prior annotations, sequence complexity, and empirical metrics on our data. Locus tiers ranged from 1 to 3, with 1 indicating the highest confidence in WGS variant detection performance, and 3 the lowest.

To specify the locus confidence tiers, we first identified regions of the genome which empirically had unusual coverage in the MGRB and 45 and Up cancer sequencing data. For each sample we defined bounds on the expected sequence coverage as the 0.001 and 0.999 quantiles of a Poisson distribution, with rate equal to the modal nonzero coverage observed across all autosomal loci within that sample. As typically 15 reads are required for high genotyping performance (Meynert et al., 2014), the lower bound was thresholded to always be at least 15. Within each sample, we defined each autosomal locus as being either in-bound (depth within the sample-specific bounds), or out-of-bound. We then calculated across all samples the rate at which each locus was out-of-bounds, considering the entire MGRB cohort. Regions for which this rate exceeded 5% (in other words, loci which had unusual coverage in at least 5% of MGRB + 45 and Up cancer samples) were marked as problematic. These problematic regions were smoothed by a morphological closing operation followed by an open operation, with structuring elements being centred intervals on the genome of size 131 bp and 11 bp, respectively, to yield a final definition of regions of unusual depth in the MGRB cohort. These regions totalled 409 Mb, 13.0% of the reference genome, 13.2% of the canonical chromosomes (1-22, X, Y), and 14.9% of the CCDS coding sequence (accessed 21 Nov 2017).

We then defined a poor-quality subset of the genome as all loci within 5 bp of the union of: the unusual depth regions, repeat regions identified by RepeatMasker, low complexity regions of the reference genome detected by mdust with default parameters, excludable regions from the ENCODE project, and poorly aligned or non-unique regions from the ENCODE project (Key Resources Table). This poor-quality subset totalled 1,832 Mb in size, 58.4% of the reference genome, 59.0% of the canonical chromosomes, and 18.1% of the CCDS coding sequence.

Variants in non-canonical chromosomes, the pseudoautosomal regions (X: 60001 - 2699520, 154931044 - 155260560; Y: 10001 - 2649520, 59034050 - 59363566), or within the poor-quality subset of the genome defined above, were assigned to the lowest confidence tier 3. For the remaining variants in canonical chromosomes, if the variant overlapped a high-confidence HG001 region identified by the GiaB consortium v3.3.2 (Zook et al., 2014) it was assigned the highest confidence tier 1, else it was assigned an intermediate confidence tier of 2. In total, 81.9% of the CCDS coding genome was in confidence tier 1 or 2 (Table 3).

**Table 3:**
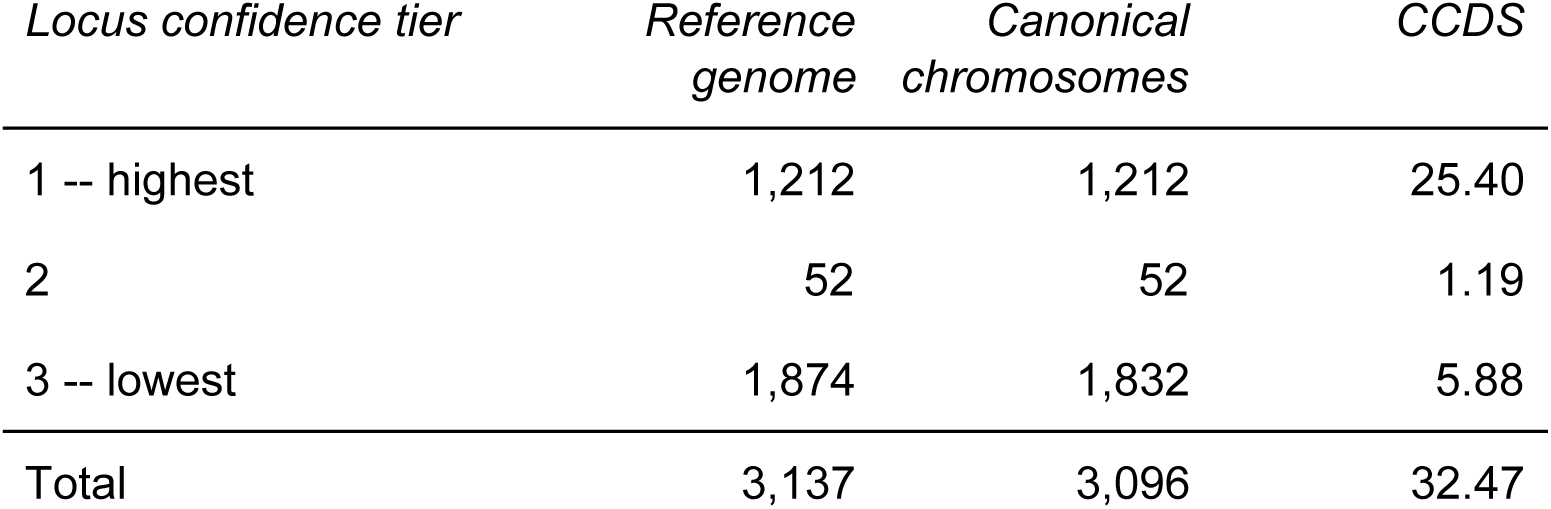
Quantity in megabases of the reference genome, the canonical chromosomes (1-22, X, Y), or the CCDS coding regions in each locus confidence tier.

#### Initial sample quality control

Poor quality MGRB and 45 and Up cancer samples were identified on the basis of genotype metrics at a small diagnostic set of loci. All 3,033 single-lane samples were genotyped at SNP loci on the Illumina Infinium QC Array 24 v1.0, using GATK GenotypeGVCFs, and quality metrics calculated within Hail v0.1 (Ganna et al., 2016). 2,904 / 3,033 (95.7%) samples passed initial quality thresholds (Table 4). Of these, 14 (0.5%) had a reported sex that did not match their genetic sex, as determined from the X chromosome inbreeding coefficient; these sex-discordant samples were not considered further. In total, 2,890 / 3,033 (95.3%) MGRB and 45 and Up cancer samples passed initial quality control (QC).

**Table 4:** Quality metric conditions for samples to pass quality control (QC). Two rounds of QC were performed, with different metric cutoffs: a first round based on genotypes at Illumina Infinium QC Array 24 SNPs only, and a second round based on genotypes called across the whole genome. Only samples passing all cutoffs in both rounds were included in the MGRB Phase 2 release.

| *Metric* | *Initial QC (Infinium SNPs)* | *Final QC (full genotypes)* |
| --- | --- | --- |
| Call rate | > 0.98 | > 0.98 |
| Depth standard deviation | < 10 | < 10 |
| VAF standard deviation at loci called heterozygous | < 1 | < 1 |
| Heterozygous :<br>Homozygous variant ratio | < 2 | < 2 |
| X chromosome inbreeding coefficient | < 0.2 or<br>> 0.8 | Not tested |
| Singleton rate | < 0.001 | Not tested |

#### Small variant genotyping, final QC

The 2,890 MGRB and 45 and Up cancer samples passing initial QC were joint called in a single batch using GATK GenotypeGVCFs, and imported to Hail v0.1 for processing. A second round of QC (Table 4) identified an additional 31 samples with poor quality metrics not revealed by the initial QC round; these were dropped. The PCRELATE component of the GENESIS 2.8.0 package (Conomos and Thornton, 2016) was used to determine structure-corrected relatedness between the 2,859 samples remaining, using autosomal SNPs LD-pruned with an r^2^ threshold of 0.1, KING robust relatedness estimates from SNPrelate 1.12.1 (Zheng et al., 2012), and without a population reference cohort. 14 pairs of individuals related to 2nd degree or closer were identified and excluded from the cohort. MGRB (cancer-free) and 45 and Up cancer samples were split into separate cohorts at this point, and four 45 and Up cancer samples excluded on the basis of incomplete or inconsistent clinical data. In summary 2,841 unrelated samples passed all data quality requirements, comprising 2,570 cancer-free MGRB individuals, 269 45 and Up cancer samples, and the reference materials RM 8391 and RM 8398.

#### Cohort population structure

The MGRB cohort population structure was determined using principal components analysis (PCA), with reference to the 1000 genomes (1000G) populations. A merged dataset of all MGRB and 45 and Up cancer genotypes and the 1000G Phase 3 genotypes (The 1000 Genomes Project Consortium, 2015) was generated in Hail. To ensure high genotype concordance between platforms, merged variants were restricted to autosomal strand-specific SNPs in Tier 1 regions of the genome (see Locus confidence tiers), with a 1000G allele frequency in the range of 5% to 95%, and no evidence of deviation from Hardy-Weinberg equilibrium within any of 17 homogeneous 1000G populations (P_HWE_ > 0.01 / 17 for each of population codes BEB, CDX, CEU, CHB, CHS, FIN, GBR, GWD, IBS, ITU, JPT, KHV, LWK, MSL, STU, TSI, and YRI). Merged variants were LD-pruned in Hail with an r^2^ threshold of 0.1, and PCA performed in Hail on biallelic variants with a combined MGRB and 1000G allele frequency in the range of 5% to 95%.

A hierarchical eigenvalue decomposition discriminant analysis classifier was constructed to assign MGRB samples to 1000G populations on the basis of PCA scores. The first classifier layer predicted a sample’s 1000G superpopulation (AFR, AMR, EAS, EUR, or SAS), and the second a sample’s European population (CEU, FIN, GBR, IBS, or TSI), conditional on EUR being the predicted superpopulation by the first layer. Models were trained on 1000G sample scores only using PC1-4 as predictors, then were applied to predict population source for the MGRB samples. All models were implemented using mclust v5.3 (Scrucca et al., 2016).

#### Small variant processing and annotation

Small variant processing and annotation was performed within Hail v0.1. Variant consequences were determined using Ensembl VEP 90 with default Ensembl release 90 databases (McLaren et al., 2016). Variants were further annotated with a range of population allele frequencies, database cross-references, and pathogenicity predictions (Table 5)

**Table 5:** Annotations applied to MGRB small variant data

| *Annotation* | *Version* | *Source or citation* |
| --- | --- | --- |
| 1000 genomes allele frequencies | Phase 3 (2 May 2013) | (The 1000 Genomes Project Consortium, 2015) |
| Haplotype reference consortium allele frequencies | 1-1 | (McCarthy et al., 2016) |
| GnomAD allele frequencies | 2.0.1 | http://gnomad.broadinstitute.org/ |
| dbSNP | 150 | (Sherry et al., 2001) |
| ClinVar | 9 Sep 2017 | (Landrum et al., 2014) |
| CATO | 1.1 | (Maurano et al., 2015) |
| Eigen coding | 1.1, 9 May 2016 | (Ionita-Laza et al., 2016) |

#### Germline structural variant detection

Germline structural variants in the MGRB and 45 and Up cancer cohorts were detected using GRIDSS v1.4.1 (Cameron et al., 2017), excluding regions in the Encode DAC Mappability Consensus Excludable list (Key Resources Table). Where possible, linked sets of breakend calls resulting from a single rearrangement were merged into higher-level structural events. To eliminate overlap with GATK indel calls and enable assessment of cohort frequencies, structural variant events were filtered to be of length at least 50 bp, and those of the same type within a window of 100 bp were merged to the one call.

Germline mobile element insertions (MEIs) were identified using Mobster v0.2.2 (Thung et al., 2014) without blacklisting existing mobile element regions. MEI calls were then processed to remove false positive events in existing mobile element regions and to estimate variant zygosity by local realignment to the reference genome. MEIs occurring in different samples within 100 bp of each other were merged to the one call.

#### Rare variant burden comparison

To compare rare variant burden between the platform-matched MGRB and 45 and Up cancer cohorts, missense or nonsense variants (as judged by VEP) in ACMG SF 2.0 cancer-associated genes were joint called across both cohorts, and each variant scored for pathogenicity by ACMG criteria, blinded to cohort. The rate of individuals carrying pathogenic variants was then directly compared by Fisher’s test. To exclude potential confounding due to source cohort, the 45 and Up component of the MGRB only was compared to the 45 and Up cancer cases.

To compare rare variant burden between the platform-mismatched MGRB and gnomAD non-Finnish European (NFE) WGS cohorts, the following procedure was used. Missense, nonsense, and synonymous variants in protein-coding genes were identified in each cohort using VEP, with identical parameters across both MGRB and gnomAD. Variants were further reduced to a very high-quality set, defined by the intersection of the Genome in a Bottle gold standard regions, and regions sequenced to a depth of at least 15 in at least 98% of samples in both the MGRB and gnomAD WGS cohorts. Variants with alternate alleles present at a frequency of 1% or greater in either cohort were discarded.

Given these high-confidence rare variant alleles, we calculated rare variant burden as 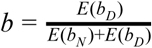, where *E*(*b*_*D*_)and *E*(*b*_*N*_)are the expected rates of individuals carrying any deleterious or all neutral genotypes in the cohort, respectively, assuming random assortment. Forms for these expectations depend on the genetic model used; for a dominant model 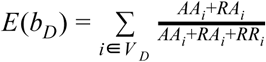, and for a recessive model 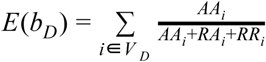, with *v*_*D*_ denoting the set of alleles called deleterious, and *AA_i_*, *RA_i_*, and *RR*_*i*_ the counts of individuals with homozygous alternate, heterozygous, and homozygous reference genotypes for allele *i*, respectively. Double-alternate heterozygous loci (eg genotype AB) were not considered in this calculation, but given the rarity of the alleles considered these were very uncommon. Expectations for neutral variation *E*(*b*_*N*_) are defined analogously, except summed over the set of neutral alleles *V* _*N*_.

To test the significance of differences in *b* between cohorts we employed a bootstrap procedure. Bootstrap draws of each source cohort genotypes were created by independent sampling with replacement of observed genotypes at each locus, and used to generate *B* = 1000 bootstrap distributions of *b*. A two-sided p-value was calculated as 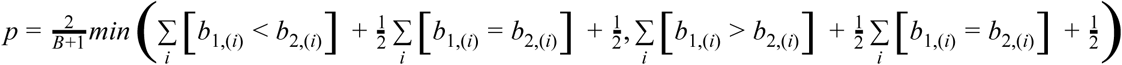, with *b*_*c,(i)*_ denoting bootstrap draw *i* of the statistic for cohort *c*.

Deleterious alleles were defined as non-synonymous changes with a CONDEL score over 0.7; neutral alleles were defined as synonymous changes. Tests examined variants in all VEP-identified protein coding genes, or variants associated with arrhythmia, cardiomyopathy, or cancer (Supplementary File 5). Eight tests were performed (two models, one neutral/deleterious classification, four gene sets).

#### Genome-wide common variant frequency comparison

To compare patterns of common variation between the MGRB and other cohorts, we merged the MGRB variants with gnomAD v2.0.1 non-Finnish European (NFE) WGS allele frequencies, and allele frequencies from a homogeneous subset of the UK BioBank genotype set, generated as previously described (Nagpal et al., 2018). To minimise the influence of technical artefacts, variants were restricted to strand-specific biallelic SNPs listed in the EBI GWAS database, that were located in regions of the genome covered by the Genome in a Bottle standard, and were sequenced to a depth of at least 15 in at least 98% of samples in both the MGRB and gnomAD WGS cohorts. Further, variants which were not observed in one or more cohorts, or were genotyped at a rate of less than 97% in any cohort, were excluded. 21,033 SNPs remained following this filtering, with very similar allele frequencies across all cohorts (Supplementary Data File 2, sheet 1; Supplementary Figure 3); these loci and frequencies were used in the following common variant analyses.

We tested for phenotype-linked bias in allele frequency between the cohorts as follows. For a given phenotype-associated set of variants, each variant was scored on two metrics: its variant allele frequency enrichment or depletion in MGRB versus gnomAD or UKBB, and the positive or negative association of the variant allele with the trait. A Fisher’s exact test was then used to test for dependence of variant enriched/depleted status on the trait direction of effect, with deviation from the null indicating an allele frequency bias between MGRB and gnomAD or UKBB that is specific to the phenotype considered.

Three sets of variants were tested by this procedure: a test set of variants reported to be associated with phenotypes depleted in the MGRB (Supplementary Data File 2, sheets 2-3), and two negative control sets of variants linked to anthropometric traits (Supplementary Data File 2, sheets 4-5), or behavioural traits (Supplementary Data File 2, sheets 6-7).

#### Polygenic score estimation and testing

Polygenic scores were calculated as 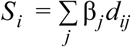 where *S*_*i*_ is the polygenic score for individual *i*, β_*j*_ the GWAS-reported coefficient for a single variant allele at locus *j*, and *d*_*ij*_ is the variant allele dosage for individual *i* at locus *j*. We considered only autosomal variants, and if a variant dosage was not available for an individual, it was imputed as 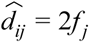, with *f*_*j*_ the variant allele frequency reported by the source publication. To reduce bias due to this imputation, variants with a call rate under 97% were excluded from polygenic score calculation in all individuals.

Polygenic score GWAS coefficients were derived from processing of summary statistics for genome-wide significant loci in reporting papers, for colorectal cancer (Schumacher et al., 2015), melanoma (Law et al., 2015), breast cancer (Michailidou et al., 2017), prostate cancer (Hoffmann et al., 2015), blood pressure (Warren et al., 2017), early-onset coronary artery disease (EOCAD) (Thériault et al., 2018), atrial fibrillation (Lubitz et al., 2017), height (Wood et al., 2014), Alzheimer’s disease (Lambert et al., 2013), and longevity (Deelen et al., 2014). GWAS coefficients were used as-is for the continuous trait of height. For all other binary traits, coefficients were converted to a log-odds scale. The Alzheimer’s disease PRS as originally reported lacked the highly significant *APOE* locus; accordingly this locus was manually added to the PRS using the tag SNP rs10414043, and an estimated β = 1.34 (Genin et al., 2011). rs10414043 was used in preference to the more conventional rs429358 as the latter was not robustly genotyped on all platforms. All loci, alleles, and coefficients used in the PRS calculations are detailed in Supplementary File 2.

An approximate bootstrap procedure was used to test for polygenic score shift between MGRB, gnomAD, UKBB, and the 45 and Up cancer cohort. All cohorts were first collapsed to allele-frequency data only, with individual genotypes discarded. For a given polygenic score and a set of cohorts to compare, testing then proceeded as follows. Polygenic score variants were first subset to those called at a rate of at least 97% in each cohort, and with an absolute difference in alternate allele frequency between MGRB, gnomAD, or UKBB of less than 4%. A bootstrap sample of genotypes was then drawn independently for each cohort and locus, polygenic scores calculated for each bootstrapped individual as above, and the mean cohort polygenic scores calculated. This was repeated for 5,000 bootstrap rounds to yield bootstrap distributions of the mean polygenic score in each cohort. To facilitate comparisons between scores of different scales, bootstrap distributions for each score were normalised by an affine transformation that brought the 0.025 and 0.975 quantiles of the MGRB scores to values of −0.5 and 0.5 respectively. The 95% bootstrap confidence intervals for each cohort were then defined as the 0.025 and 0.975 quantiles of the normalised scores. Approximate two-sided p values for the overlap between two bootstrap distributions were estimated as 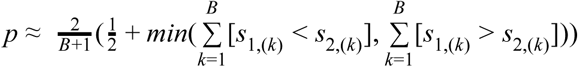 where *s*_*1,(k)*_ and *s*_*2,(k)*_ are samples from the the bootstrap mean values for cohorts 1 and 2, respectively, *B* = 100000 is the number of samples taken with replacement from the bootstrap mean values, and [ ] denotes Iverson brackets.

The statistical power improvement from using the MGRB extreme phenotype cohort as a control versus gnomAD was estimated by asymptotic approximation. Bootstrap distribution means of the mean prostate cancer score difference between the 45 and Up cancer cases, and gnomAD and MGRB controls, were used as mean shift values for statistical power calculation. After correcting for varying cohort sample size, bootstrap distribution variance was highly consistent across all three cohorts, and pooled variance scaled to a sample size of 1 was used as the dispersion parameter. Power was then calculated across a range of sample sizes for both the MGRB vs 45 and Up cancer, and gnomAD vs 45 and Up cancer tests, by direct root finding of the relevant t distributions. Finally, the power vs sample size relationship was inverted by piecewise linear interpolation to yield the sample size vs power curves.

Individual genotypes were available for both MGRB and 45 and Up cancer cohorts. In these cases, a secondary analysis was performed that directly compared the distributions of individual polygenic scores between cohorts. Height prediction was validated by ordinary linear regression of measured individual height against the polygenic height predictor (Wood et al., 2014) with additional additive linear covariates of sex and age at measurement; no evidence for model misspecification was observed. The association between polygenic score and risk of specific cancers was assessed by logistic regression, with the effect of polygenic score on cancer risk modelled by GCV-penalised thin plate splines. Comparisons were restricted to the specific cancers of prostate, colorectal, and melanoma, as other cancers were either poorly sampled in the 45 and Up cancer cases, or did not have polygenic scores defined.

#### Incidental somatic variant detection

Somatic variants were identified in post-BQSR BAM files using FreeBayes, with options:--pooled-continuous --standard-filters --min-alternate-fraction 0 --min-alternate-count 3 --hwe-priors-off --allele-balance-priors-off --use-mapping-quality. FreeBayes was restricted to detecting variants within 10 kb of RefSeq genes in the COSMIC Cancer Gene Census (Forbes et al., 2017) downloaded 11 December 2017. Variant annotation was performed using the Ensembl VEP (McLaren et al., 2016) release 90, with default options, and variants were notated with COSMIC 83 frequencies.

Annotated variants were filtered to retain only non-synonymous variation (missense, splice donor or acceptor, start lost, stop gained, frameshift, or inframe indel) affecting Cancer Gene Census Tier 1 genes, with a maximum population allele frequency of less than 0.1%, a variant allele fraction (VAF) of at least 10%, and three or more reads supporting the variant. We then identified likely driver mutations from these filtered variants by the following criteria: either a variant had a HIGH consequence in a canonical tumour suppressor gene transcript, or the variant was observed at least 100 times in the COSMIC database. Consequences and canonical transcripts were as defined by Ensembl VEP; tumour suppressor genes were Tier 1 genes from the COSMIC Cancer Gene Census with a TSG annotation.

#### Sequence-based measures of age

##### Telomere length

Telomere lengths were estimated using Telseq v0.0.1 (Ding et al., 2014). To reduce batch effects between the ASRB and MGRB cohorts, ASRB telomere length estimates were calibrated using Deming regression, fit to 85 ASRB samples sequenced both in the original ASRB batch, and contemporaneously with the MGRB.

Telomere length estimation by Telseq was validated by qPCR on a subset of 120 samples from the ASRB and MGRB cohorts (Supplementary Figure 7), as described previously (Cawthon, 2002) with minor modifications. Briefly, qPCR was conducted in triplicate.

Reactions included: genomic DNA (5 ng), 2x Rotor-Gene SYBR Green Master Mix (Qiagen), 500 nM Tel forward [5’-CGGTTT(GTTTGG)_5_GTT-3’] and 500 nM Tel reverse

[5’-GGCTTG(CCTTAC)_5_CCT-3’] or 300 nM 36B4 forward

[5’-CAGCAAGTGGGAAGGTGTAATCC-3’] and 500 nM 36B4 reverse

[5’-CCCATTCTATCATCAACGGGTACAA-3’] in a 25 μL reaction. Amplification was conducted in a Rotor-Gene Q qPCR cycler (Qiagen) at 95°C for 5 min, followed by 30 cycles of 95°C for 7 sec and 58°C for 10 sec (telomere reaction) or 35 cycles of 95°C for 15 sec and 58°C for 30 sec (single copy gene reaction). Telomere content for each sample was determined by the telomere to single copy gene ratio (T/S ratio) by calculating ΔCt (Ct_telomere_/ Ct_single copy gene_). The T/S ratio of each sample was normalized to the mean T/S ratio of a reference sample, which was included in each run. The experiment was accepted if the reference sample T/S ratio ranged within 95% variation interval, and if the standard curve had a high correlation factor (R^2^ > 0.95).

##### Mitochondria and Y chromosome copy number

Mean mitochondrial genome copy number in each sample was estimated using read counts, as 2 × (*R*_*MT*_÷ *S*_*MT*_) ÷ (*R*_*A*_ ÷ *S*_*A*_), where *R*_*Z*_ and *S*_*Z*_ denote the number of reads mapping to contig set *Z* and the total size of contig set *Z*, and *MT* and *A* denote mitochondrial and autosomal contigs, respectively. Read counts were mapped and aligned reads reported by samtools idxstats, and were not corrected for read duplication. Patch contigs were not included in counts. Y copy number in males was estimated by an analogous procedure, as 2 × (*R_Y_*÷ *S_Y_*) ÷ (*R_A_*÷ *S_A_*).

##### Mitochondrial variants

Variants in the mitochondrial genome were detected using FreeBayes, considering only reads with base quality over 24 and mapping quality over 30; all other parameters were left at defaults. Variants with fewer than 10 alternate reads, or an alternate allele fraction under 0.001, were discarded. For each variant passing these filters a Phred-like quality score *q* was calculated as *q* =− 10*log*_10_(1 − *F* (*n*; *p*, *N*)), with *n* the count of alternate allele reads, *N* the total depth at the variant locus, *p* = 0.0025 a fixed error rate estimate, and *F* (*n*; *p*, *N*) the cumulative density function of a binomial distribution with *N* draws and success probability *p*. Variants with *q* < 30, high depth variants (*n* > 15) with an alternate read strand bias of greater than 0.9, or variants in the highly variable locations MT:302-319 or MT:3105-3109 were discarded. The final metric of mitochondrial variant burden for a sample was defined as the number of low-frequency (variant allele fraction under 0.01) variants passing all above filters in that sample.

##### Somatic single nucleotide variants

Somatic SNV burden was estimated using a combination of statistical filtering and spectral denoising. Putative somatic SNVs were first identified on the basis of a variant allele frequency that was statistically inconsistent with either machine error or germline variation. The burden of these variants in each sample was then dimensionally reduced by spectral factorization, and per-sample signature scores used as the final somatic variant estimates.

We first developed a statistical filtering procedure to identify likely somatic variants, that uses dynamic thresholds to optimise sensitivity while controlling signal to noise ratio. This procedure calls a variant at a given locus as likely somatic if it satisfies the following criterion:

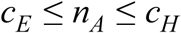

where *n*_*A*_ is the number of non-reference allele reads at the locus, and *c*_*E*_ and *c*_*H*_ are integers which maximise:

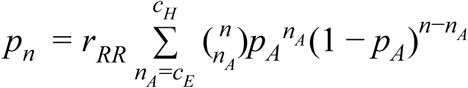

subject to:

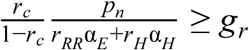

with 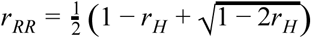 and 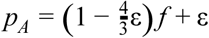. Here *n* is the sum of reference and alternate allele depths at the locus, *r*_*H*_ is the expected rate of heterozygous variant germline loci, *r*_*C*_ the expected rate of somatic variant loci, ε the base read error rate, *g*_*r*_ the minimum acceptable ratio of true positive calls to false positive, and *f* is the expected somatic variant allele fraction. α_*E*_ and α_*H*_ are test sizes corresponding to thresholds *c*_*E*_ and 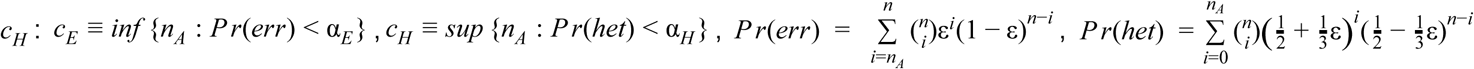. Informally, this procedure selects variants with an alternate allele count *n*_*A*_ too large to be due to sequencing error (*n*_*A*_ ≥ *c*_*E*_), yet too low to be from a poorly-sampled heterozygous germline locus (*n*_*A*_ ≤ *c_H_*). The derivation of this procedure and further details are available in Supplementary File 4. A notable advantage of this procedure is that it yields per-locus estimates of variant detection sensitivity, as *p*_*n*_. These estimates are critical in normalization of variant detection rates to account for differential coverage across variable sequencing runs, which is necessary for the accurate estimation of sample somatic variant burden. Insufficiencies of the simple error model used above result in incomplete control of the signal to noise ratio, and further filtering is required to reliably quantify somatic variant burden. Assuming that true somatic events and false positive machine noise exhibit differential sequence context bias, we use a spectral dimensionality reduction approach to achieve additional denoising and summarise the total somatic variant burden in each sample. Extending previous cancer somatic signature work (Alexandrov et al., 2013; Gehring et al., 2015), we calculate per-sample sensitivity-normalised somatic variant burden for each of 96 single-nucleotide variant classes, as the total number of detected somatic events of a given class in that sample, divided by the sum of *p*_*n*_ in that sample for all loci corresponding to the given variant class. The resulting 96 × *n* normalised burden matrix is then reduced by non-negative matrix factorization (Brunet et al., 2004), using 100 random optimisation starting points per cardinality. To select the appropriate factorization cardinality, we reduce the burden matrix by merging groups of 16 age-consecutive samples by summing burden for each variant class, and factorize this reduced matrix with 100 random restarts and cardinality ranging from 2 to 10. The lowest cardinality that gives an inflection point on the plot of explained variance versus cardinality is selected, and applied to the full burden matrix. Per-sample scores are extracted from the best of 100 random runs at this final selected cardinality.

We applied the above procedure to the MGRB and ASRB samples, with parameters adapted to maximise sensitivity with low-depth sequencing data: *g*_*r*_ = 5, *f* = 0.2, ε = 2.0 × 10^−3^, *r*_*H*_ = 1.0 × 10^−4^, *r*_*C*_ = 5.0 × 10^−7^. Our filtering process employed SNVs identified by samtools mpileup, with maximum depth 101, mapping quality adjustment of 50, BAQ recalculation, no indel reporting, and minimum read and mapping qualities of 30, and employed a blacklist of common SNPs observed in either MGRB or dbSNP. The factorization cardinality procedure applied to our data indicated that three signatures best described the mutation patterns observed (Supplementary Figure 8). Signature 3 in this work was quantitatively similar to COSMIC signature 5 (cosine similarity 0.81), previously reported to be associated with age at cancer diagnosis (Alexandrov et al., 2015), and the per-sample scores for this signature were used as the summative somatic burden measure. Signature 1 from this work was very similar to COSMIC signature 1 (cosine similarity 0.95), which has also been associated with spontaneous deamination processes and age. However, we observed substantial inter-cohort differences in score distribution for this signature, suggestive of high technical variability, and did not examine it further.

##### Somatic copy number variants

We developed a model-based strategy to identify subclonal copy number variants (CNVs), assuming a single genetically homogeneous subclone present on a background of diploid cells.

We first defined a set of autosomal SNPs with highly stable sequencing characteristics on our platform. We selected loci containing autosomal biallelic SNPs in the MGRB cohort, with a variant allele fraction between 5% and 95%, and a mean GC content in the surrounding 100 bp of between 30% and 55%. These were further filtered to retain only loci with highly consistent coverage in both the MGRB and ASRB cohort data, with mean(DP_rel_) ∈ [0.9, 1.1] in both cohorts, var(DP_rel_) ∈ [0.025, 0.033] in the MGRB, and var(DP_rel_) ∈ [0.025, 0.040] in the ASRB cohort. Here DP_rel_ is locus depth relative to mean sample depth, and statistics are calculated over all samples in each cohort. 1,862,065 loci passed all filters, with a median inter-locus distance of 626 bp, and 5th and 95th percentiles of 30 and 4,904 bp, respectively.

We individually genotyped MGRB, 45 and Up cancer, and ASRB samples at this set of highly reliable loci using GATK HaplotypeCaller with default parameters, except for a variant window size of 100 bp (-ip 100). Within each sample, the depths of reference and variant alleles at all heterozygous SNV target loci were fit to the following subclonal CNV model, to produce estimates of local ploidy and global sample subclonal fraction.

Consider a locus *i* in a single sample which contains fraction *f* of aneuploid cells, the remaining 1 − *f* being entirely diploid (gonosomes are not modeled). We denote the copy number (ploidy) of the aneuploid cells at *i* with *k*_1*i*_ and *k*_2*i*_, 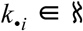. For example, *k*_1*i*_, *k*_2*i*_ = (1, 1) denotes a diploid state (no aneuploidy), *k*_1*i*_, *k*_2*i*_ = (1, 0) the deletion of one allele, and *k*_1*i*_, *k*_2*i*_ = (2, 2) duplication of both alleles. Our task is to estimate *k*_1*i*_, *k*_2_*_i_* for all *i*, and *f* globally, given reference and non-reference allele depths *d*_*ri*_ and *d*_*ai*_.

The extent to which the aneuploid cell ploidies *k*_1*i*_ and *k*_2*i*_ affect the representation of alleles in the mixed cell population depends on the aneuploid cell fraction *f*. Let *p*_1*i*_ and *p*_2*i*_ represent the mean ploidy of each chromatid in the mixed cell DNA pool. As the pool is assumed to consist of only two populations, with 1 − *f* of the cells diploid, *p*_1*i*_ = *f k*_1*i*_ + (1 − *f*) and *p*_2*i*_ = *f k*_2*i*_ + (1 − *f*).

We assume that the sequencer does not exhibit allelic bias. Then, *E*[*d*_1*i*_] = *c_i_p*_1*i*_, *E*[*d*_2*i*_] = *c_i_p*_2*i*_, with *c_i_*a normalising constant to account for the depth at locus *i*. Here *d*_1*i*_ and *d*_2*i*_ denote the depths of reads from chromatid 1 and 2, respectively. Unfortunately we do not have phased genotypes, and so cannot easily determine the chromatid source of each read. Instead we have unphased reference and non-reference depths *d*_*ri*_ and *d*_*ai*_ must account for the resulting phase uncertainty with a mixture model.

Disregarding allele phasing we model the depths of reference and non-reference reads at i using a mixture: *d_ri_*, *d_ai_*∼ *D*(*c_i_p*_1*i*_), *D*(*c_i_p*_2*i*_) with probability 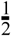, else *d*_*ri*_, *d*_*ai*_ ∼ *D*(*c*_*i*_p_2*i*_), *D*(*c_i_p*_1*i*_), *D*(μ) denoting a distribution function with expected value μ. In our implementation we employ a negative binomial distribution for *D*, with probability mass function 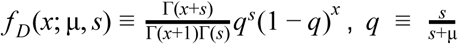. The size term *s* captures overdispersion relative to the Poisson distribution, and is optimised per-sample in the model fit.

The normalising constant *c*_*i*_ is half the expected depth at locus *i*, which is itself a complex function of locus- and sample-specific factors. We model this function at the locus- and sample-level using empirical cohort depth measurements, and a sample-specific GC bias correction. Specifically, we define *c*_*i*_ ≡ *b*_*i*_*exp*(*h*(*g*_*i*_)), where *b*_*i*_ is the mean relative depth of locus *i* (where relative depth is defined as 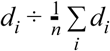, with *d_i_*≡ *d_ri_*+ *d_ai_*and *n* the number of target loci, *n* = 1862065), across the sample’s cohort (either MGRB or ASRB), *h* is a smooth function, and *g*_*i*_ is a vector of GC fraction in windows of various size around locus *i*. For this work, *g*_*i*_ was a 5-vector of GC fraction in windows of size 100, 200, 400, 600, and 800 bp, calculated on the reference sequence centered at locus *i*. The sample-specific GC correction function *h* was implemented using a generalised additive model (GAM) with five smooth terms, and fit to all heterozygous loci for each sample as *ln*(*c*_*i*_ ÷ *b*_*i*_) ∼ *s*(*r*_1*i*_) + *s*(*r*_2*i*_) + *s*(*r*_3*i*_) + *s*(*r*_4*i*_) + *s*(*r*_5*i*_), with *r*_*ji*_ being the score of the *j* th principal component of the GC fraction matrix for locus *i*, and *s* denoting a penalised regression spline term. GAMs were fit using mgcv 1.8-17 (Wood, 2004) with default parameters.

A greedy agglomerative algorithm was used to segment the genome of each sample into regions of differing ploidy state. Initially the genome was divided into segments of 100 consecutive heterozygous loci, with segment boundaries enforced between chromosomes. Adjacent segments were tested for identical distribution of *d*_*ri*_ ÷ *c*_*i*_, *d*_*ai*_ ÷ *c*_*i*_ by a 2-sample Kolmogorov-Smirnov test, and the two segments with the highest p value genome-wide were merged. This process was repeated until either no segment pairs remained to merge, or all Kolmogorov-Smirnov test p values were less than 0.01. Segments were never merged between chromosomes.

The above model was fit to the allele counts within each genome segment by maximum likelihood. Ploidies of each segment, *k*_1*i*_ and *k*_2*i*_, as well as the global aneuploid fraction *f* and overdispersion *s*, were optimised by grid search with local polishing. As very high ploidies coupled with low *f* result in highly expressive but likely incorrect models, the maximum allowable ploidy *k_max_*; *k*_1*i*_, *k*_2*i*_ ≤ *k*_*max*_was determined in an outer loop through minimisation of the Bayesian Information Criterion (BIC). A final polishing step was applied to the BIC-optimal model, which merged consecutive segments of the genome if they were assigned identical chromatid ploidies by the model. This final polished model yielded global cell fraction *f*, as well as local ploidies across the genome, for the single aneuploid clone assumed to be present in each sample.

##### Clonal haematopoiesis

Extending previous work (Jaiswal et al., 2014), clonal haematopoiesis of indeterminate potential (CHIP) was defined in an individual if either: a somatic small variant (see section *Incidental somatic variant detection*) was detected with a variant allele frequency of at least 10%, or somatic copy number variation (see section *Somatic copy number variants*) indicated the presence of a clone comprising at least 10% of nucleated blood cells.

##### Somatic burden statistical analysis

Exploratory analysis indicated that variable transformation was required for some measures. For the following analyses, telomere length and Y copy number were modelled as-is; somatic variant burden, mitochondrial load, and mitochondrial variant count were log-transformed prior to modelling; and grip strength in kg was power transformed with exponent 0.7.

Within-cohort trends in somatic measures were estimated by linear regression, with 95% Wald confidence intervals. Likelihood ratio tests of nested models were used to evaluate inter-cohort trend differences, with p-values corrected for multiple testing by Holm’s step-up procedure (Holm, 1979).

We used a permutation procedure to test the importance of somatic burden measures in predicting grip strength and gait speed, conditioned on age. For each of eighteen possible frailty measure x somatic measure x sex combinations (Frailty measures: grip strength, gait speed; Somatic measures: Telseq telomere length, nuclear somatic variant burden, mtDNA copy number, mitochondrial variant count, and Y copy number in males only), we calculated the deviance of the following generalised additive model *frailty* ∼ *s*(*age*) + *s*(*weight*) + *s*(*BM I*) + *s*(*abdocirc*) + *s*(*somatic*), with age in years, weight in kg, BMI in kg/m^2^, abdominal circumference (abdocirc) in cm, and the somatic measure of interest (transformed if relevant following exploratory analysis). In this model specification, *s*(*x*) denotes a GCV-penalised thin plate spline smooth term in *x* as implemented in R package mgcv (Wood, 2004), with Gaussian error and identity link. This model’s deviance *d* was compared to the deviance *d*_(*i*)_ of 10,000 models fit in the same manner but with the somatic variable permuted, and a p-value estimated as 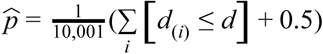. To address multiple testing concerns we used a two stage process. In the first stage p-values were calculated as above for all 18 tests on a randomly selected subset of 25% of the ASPREE samples. Tests with a p-value less than 0.2 in the first stage were tested in the second validation stage on the remaining 75% of the ASPREE samples, and these second-stage p-values corrected for multiple testing by Holm’s method.

We observed cohort differences in intercepts in plots of somatic measures versus age. To remove these solely for the purposes of illustration (Figure 3), for each somatic measure we fit the generalised additive model *measure* ∼ *s*(*age*, *by* = *sex*) + *cohort*, with Gaussian error and identity link. In this model specification *s*(*age*, *by* = *sex*) denotes a GCV-penalised thin plate spline with age as the predictor variable, stratified by sex. Model fits were performed using the R package mgcv (Wood, 2004). After confirming the suitability of the model fits, cohort-specific effects were removed by calculating the quantity 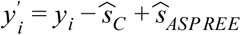 for each individual and measure, where 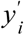 is the cohort-corrected somatic measure for individual *i*, to be plotted; *y*_*i*_ is the original measurement for individual *i* in cohort *C*; and 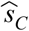 and 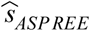 are the model estimates of the cohort intercept term for cohort *C* and the ASPREE cohort, respectively. In this manner, somatic measurements were transformed to have an intercept matching that fitted to the ASPREE cohort.

We used the following procedure to illustrate the effect of mtDNA copy number on grip strength in males. For each male individual *i* in the ASPREE cohort, an age-local quantile of mitochondrial DNA copy number *c*_*i*_ was defined as 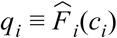, where 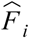 is the empirical cumulative distribution function of *c* in the neighbourhood of individual *i*, with the neighbourhood of an individual *i* defined as all male ASPREE individuals within ± 1 year of age of *i*. Ages were rounded to the nearest integer for the purposes of neighbourhood definition; for the median ASPREE male age of 80 years, this neighbourhood contained 293 men with ages in [79, 81] years. Given these local mtDNA copy number quantile estimates *q*, a generalised additive model of the form *gripstrength* ∼ *age* + *s*(*q*) was fit using the R package mgcv (Wood, 2004), with *s* smooth term as above. Predictions from this model with *age* = 80 and varying *q* defined the estimated influence of age-local mtDNA copy number on grip strength for an 80 year old man. These grip strength predictions were transformed to effective age estimates assuming typical mtDNA copy number by inversion of the model predictions for *s* = 0.5, and used to calculate an age excess as a function of *q*. Variability of this relationship was estimated using 100,000 bootstrap samples, and results presented as highest posterior density intervals.

## Data and Software Availability

Summary variant frequency data for the MGRB cohort are available at the web portal: https://sgc.garvan.org.au/explore. Complete genotype, phenotype, and raw data are available upon application to Prof. David Thomas, or. Source code for all analyses is available at https://github.com/mpinese/mgrb-manuscript; source code for the somatic SNV and LoH detection tools can be found at https://github.com/mpinese/soma-snv and https://github.com/mpinese/soma-cnv.

## Additional Resources

The Medical Genome Reference Bank web portal: https://sgc.garvan.org.au/explore

## Key Resources Table

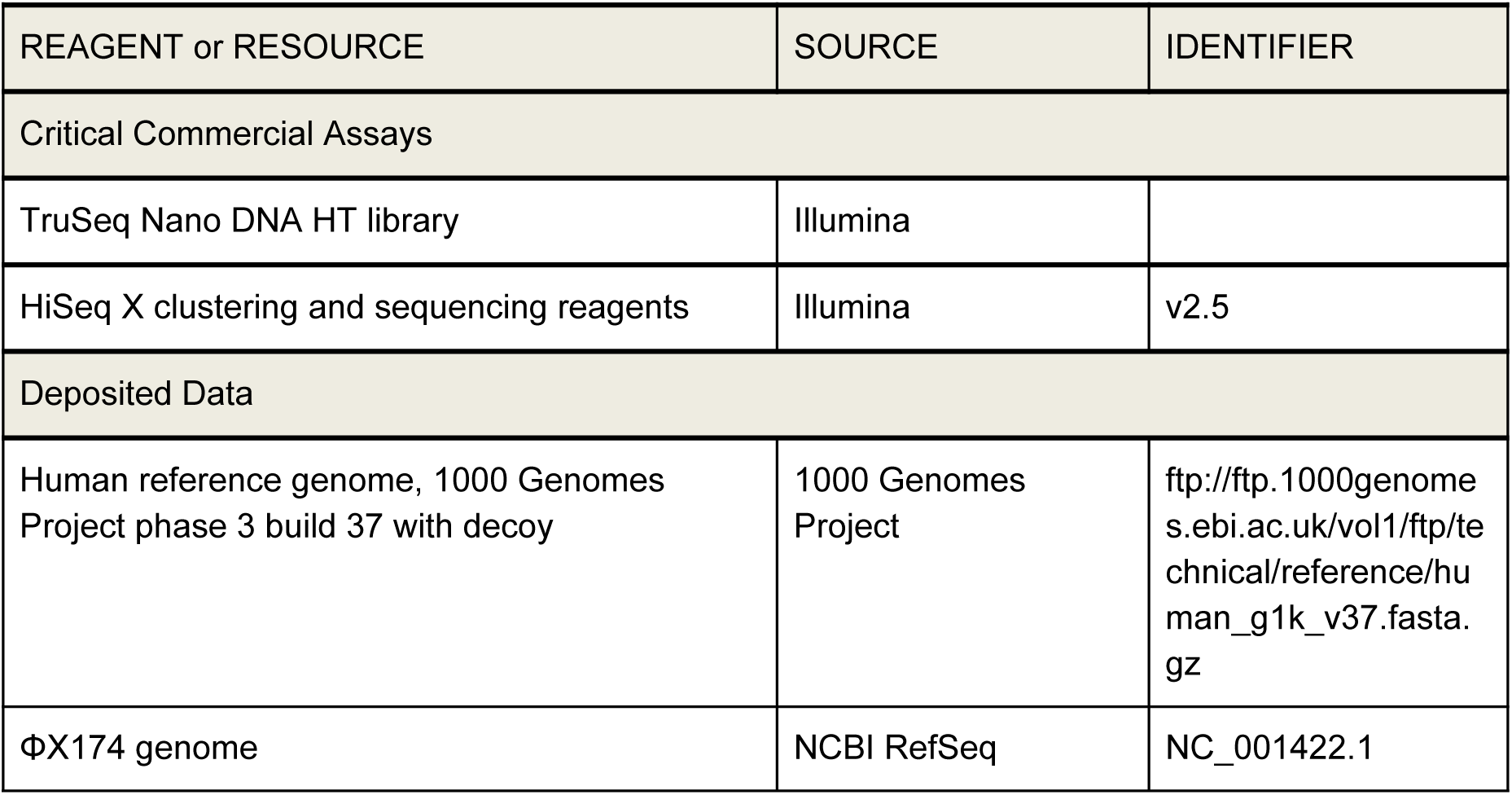

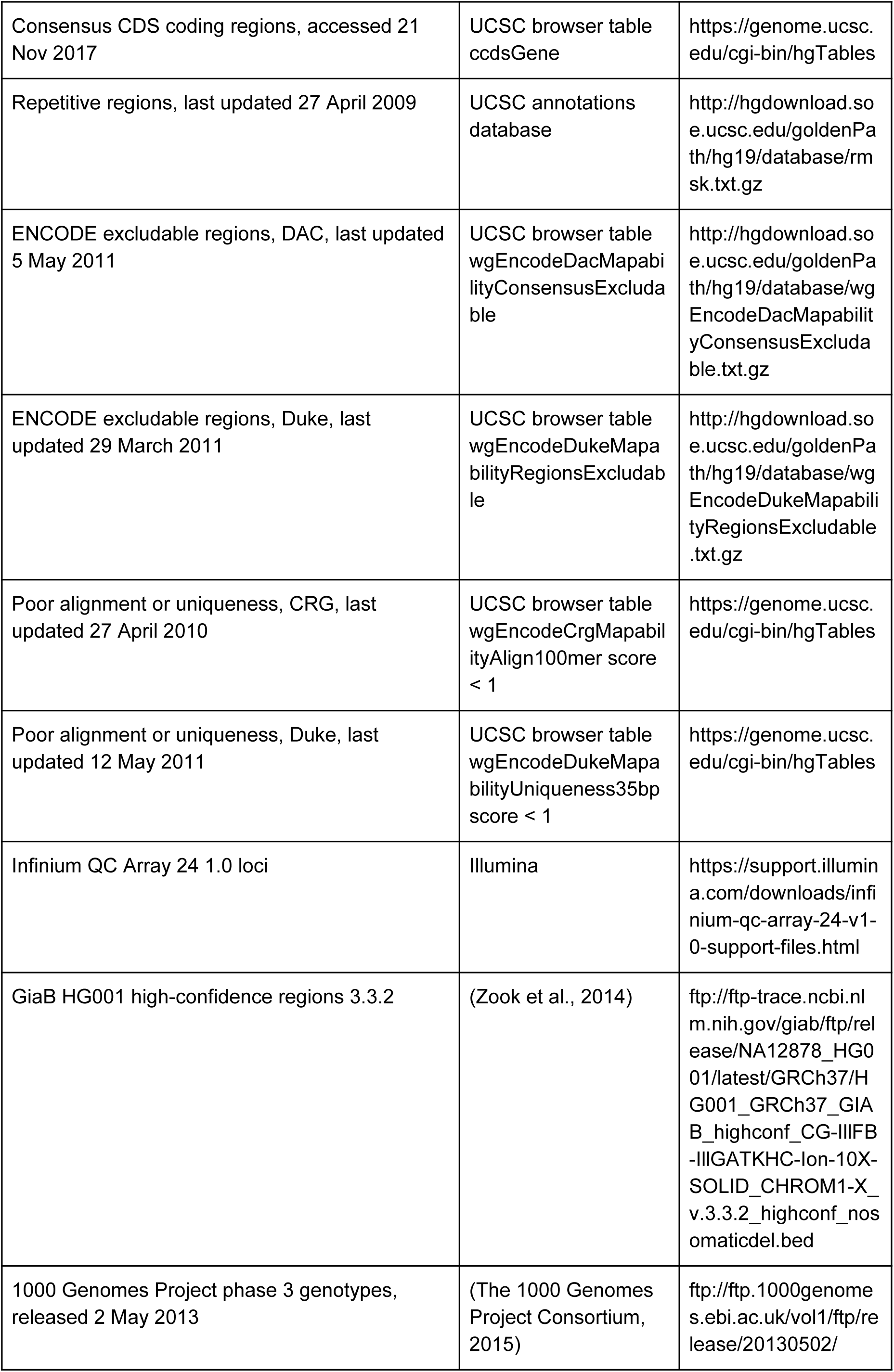

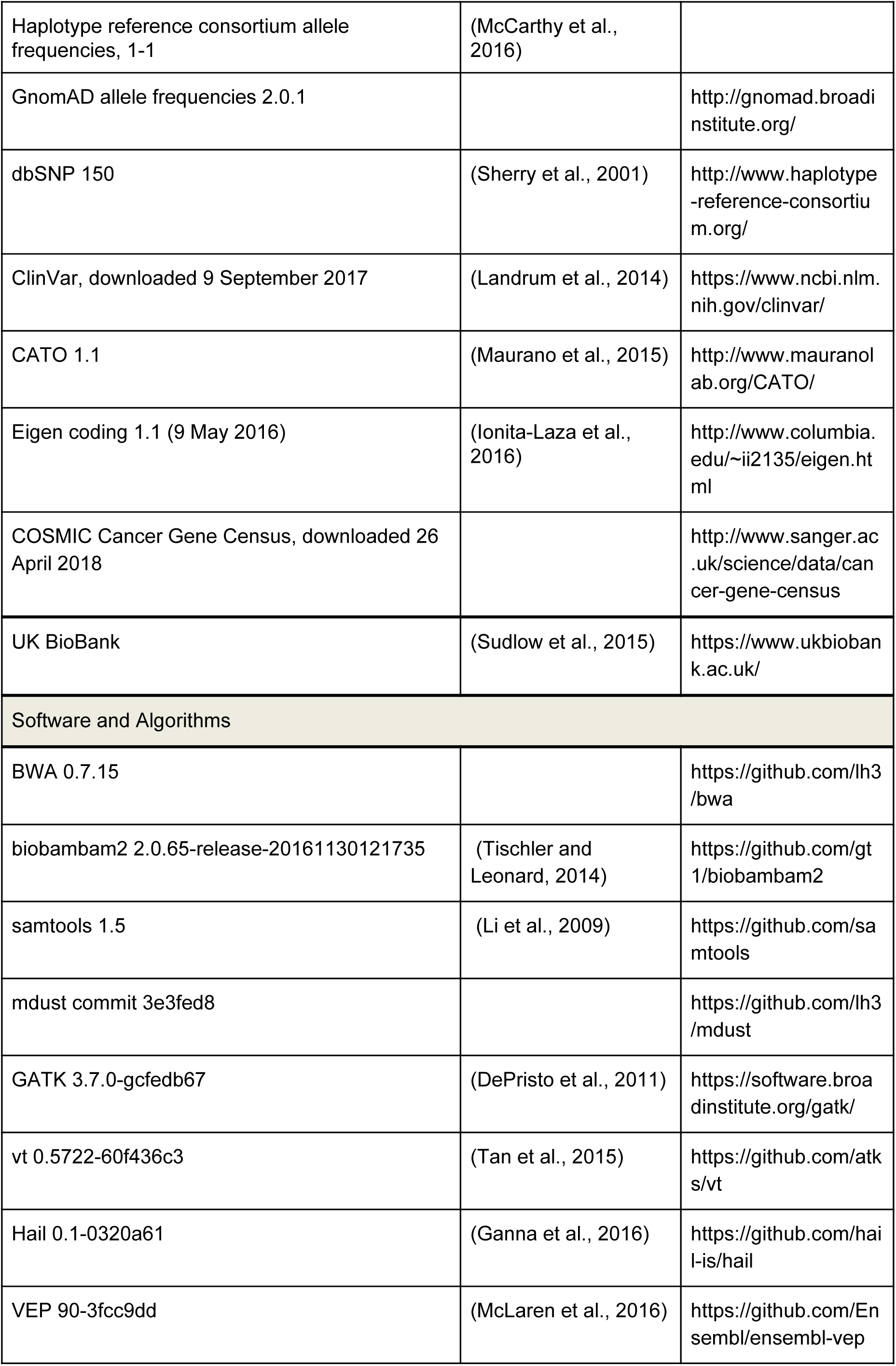

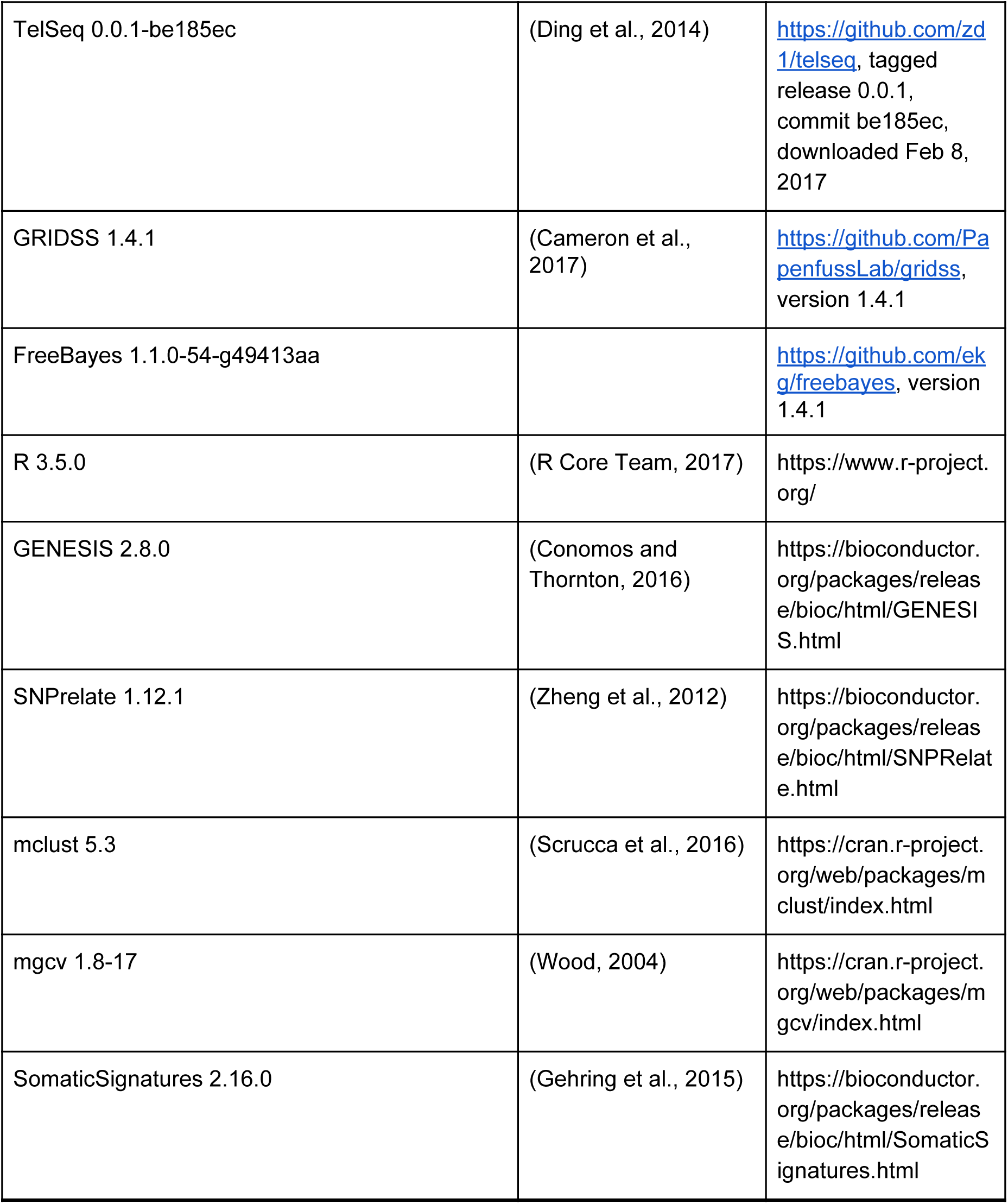

## Supplemental Information

**Supplementary Figure 1:**
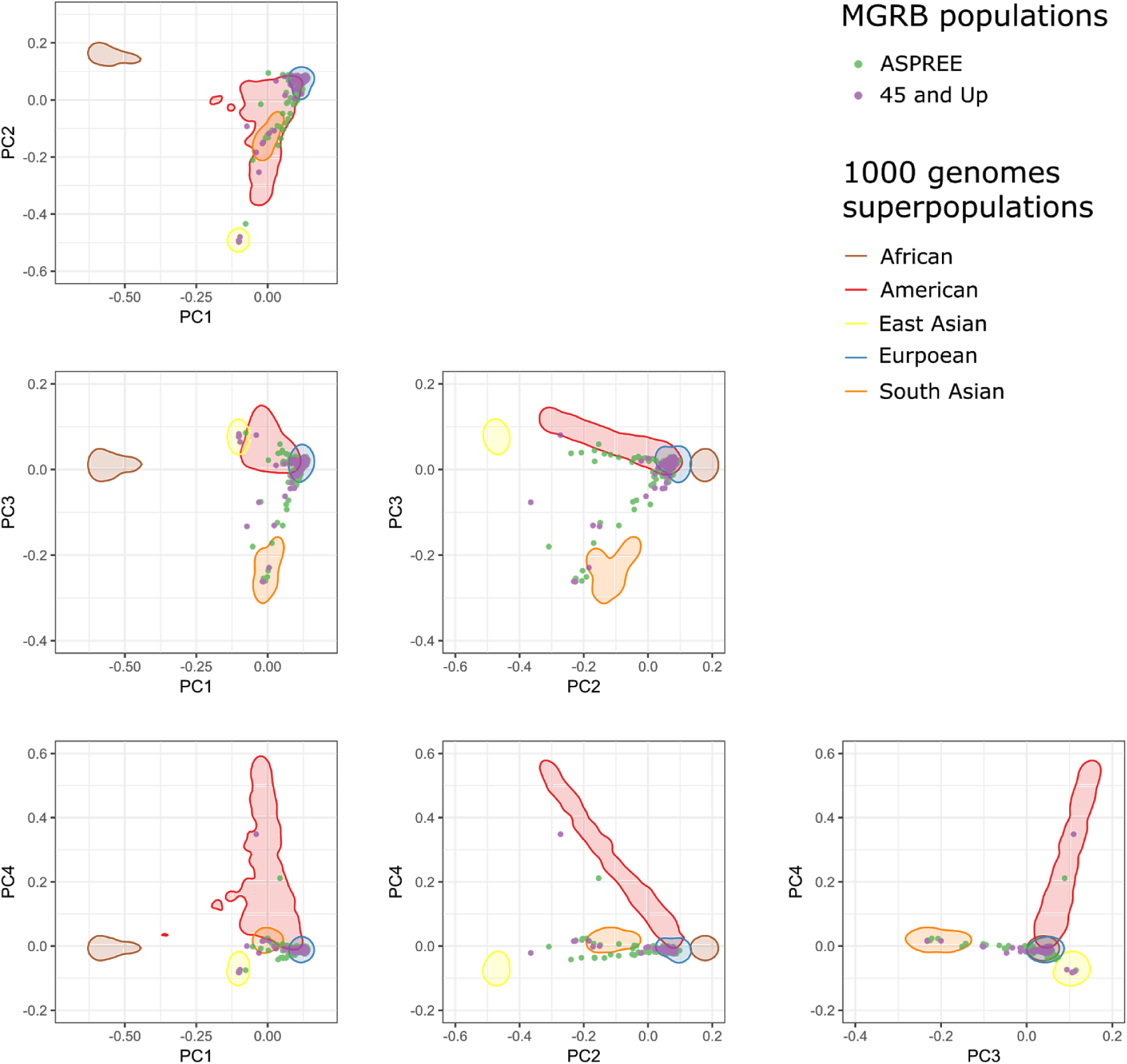
Population structure in the MGRB. The MGRB was combined with the 1000 Genomes cohort at high-confidence SNVs, and PCA was performed following LD pruning. Four strong components resulted, scores for which are shown relative to 95% kernel density estimates of the 1000 Genomes superpopulations. The MGRB cohort was largely homogeneous and clustered with the 1000 Genomes European superpopulation.

**Supplementary Figure 2:**
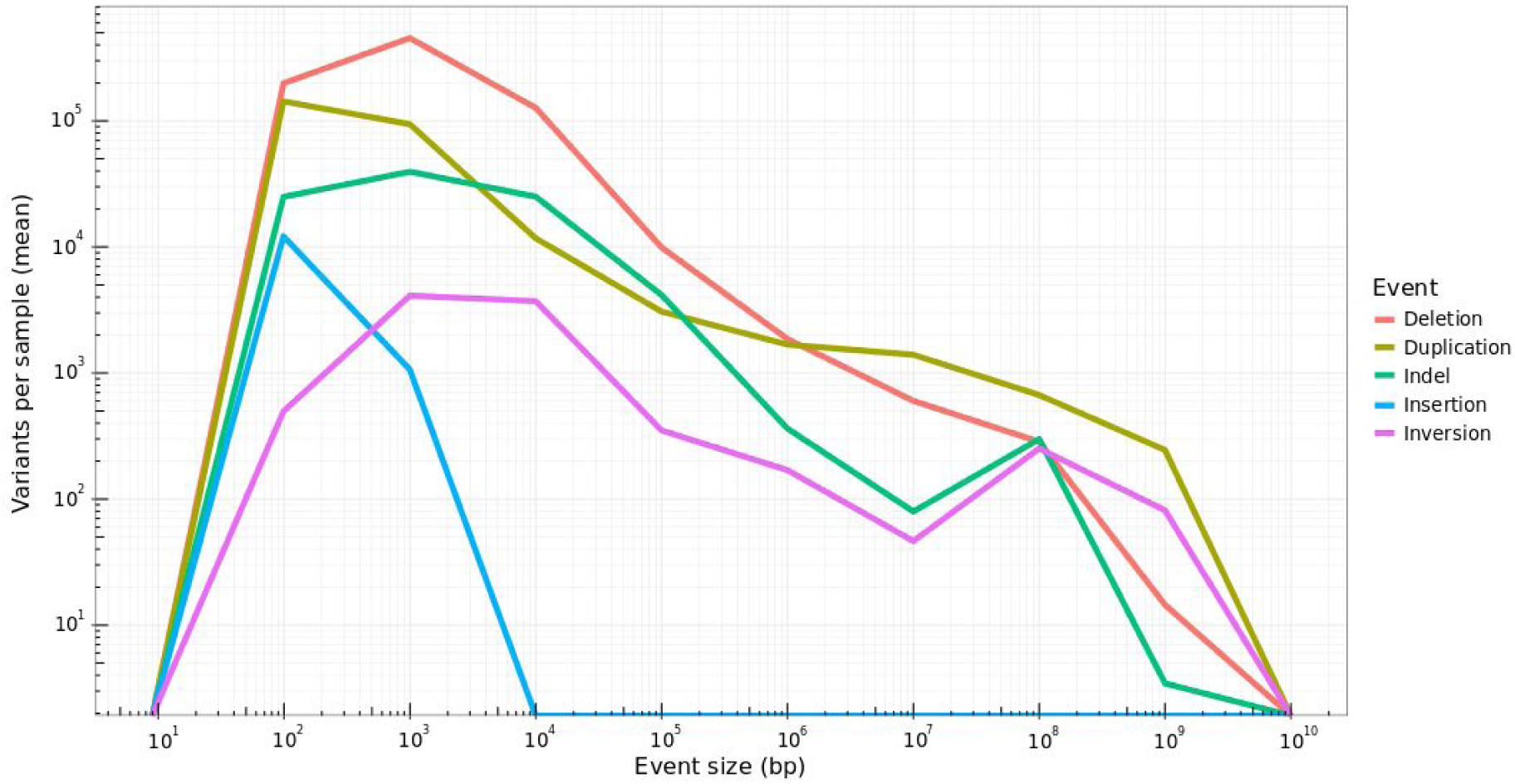
Distribution of structural variant event types and sizes detected in the MGRB by GRIDSS.

**Supplementary Figure 3:**
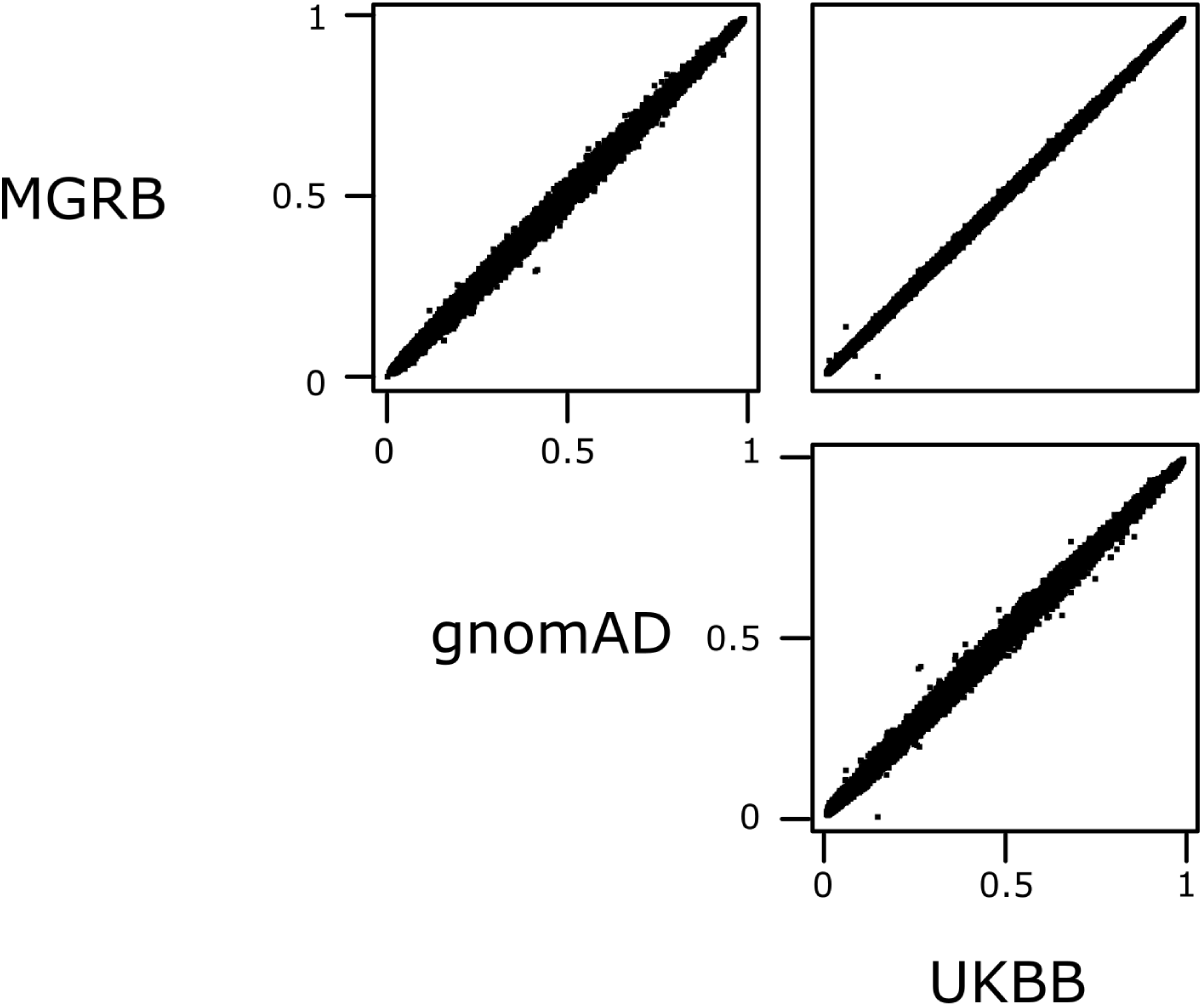
SNP alternate allele frequencies compared between MGRB, gnomAD, and UK BioBank cohorts. Strand-specific biallelic SNPs in well-called regions and reported in the EBI GWAS catalogue only shown.

**Supplementary Figure 4:**
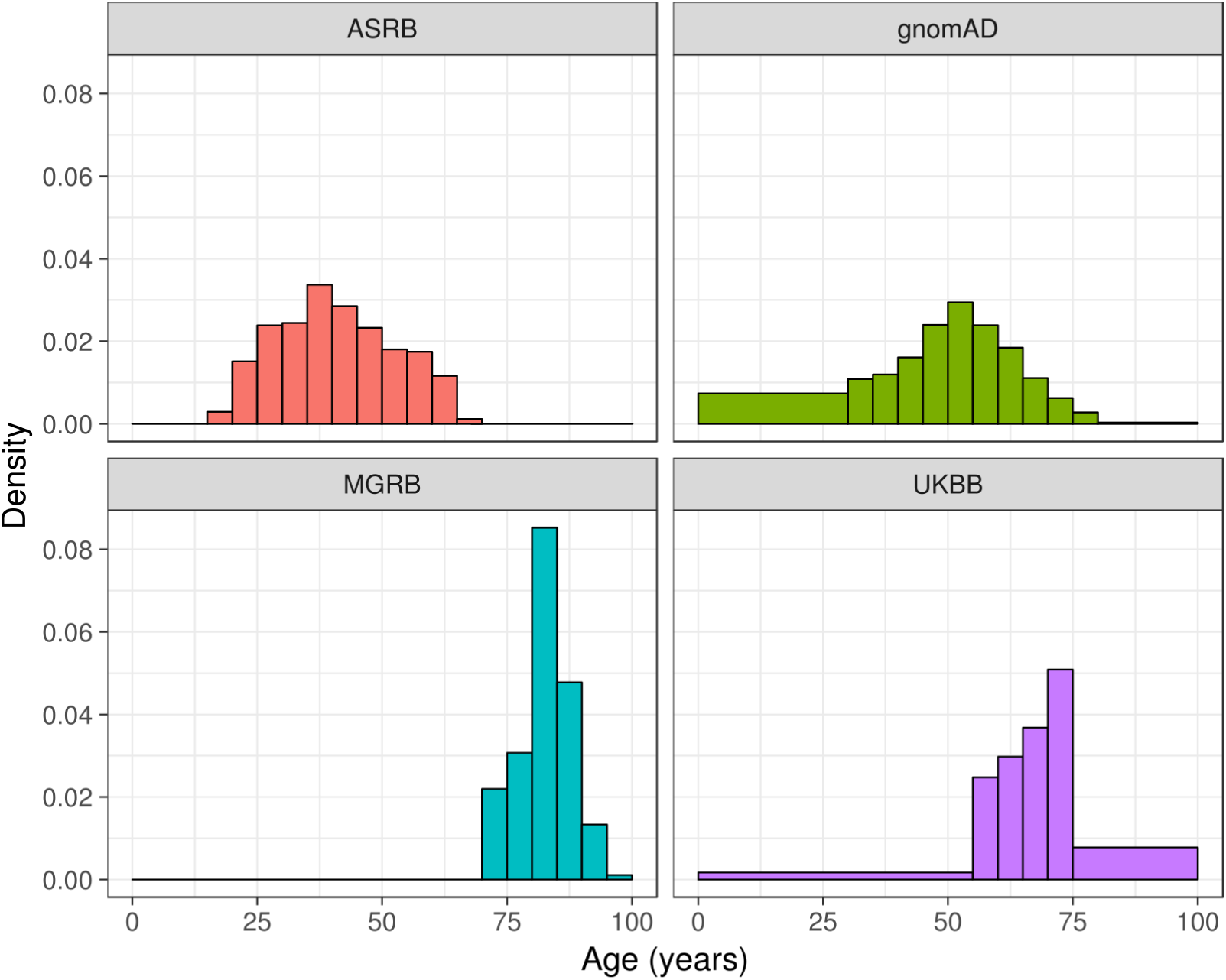
Distribution of participant ages in the Australian Schizophrenia Research Bank (ASRB), gnomAD, MGRB, and UK BioBank (UKBB) cohorts. Ages were truncated at 100 years and binned into five year intervals except for terminal bins, which vary in size as shown.

**Supplementary Figure 5:**
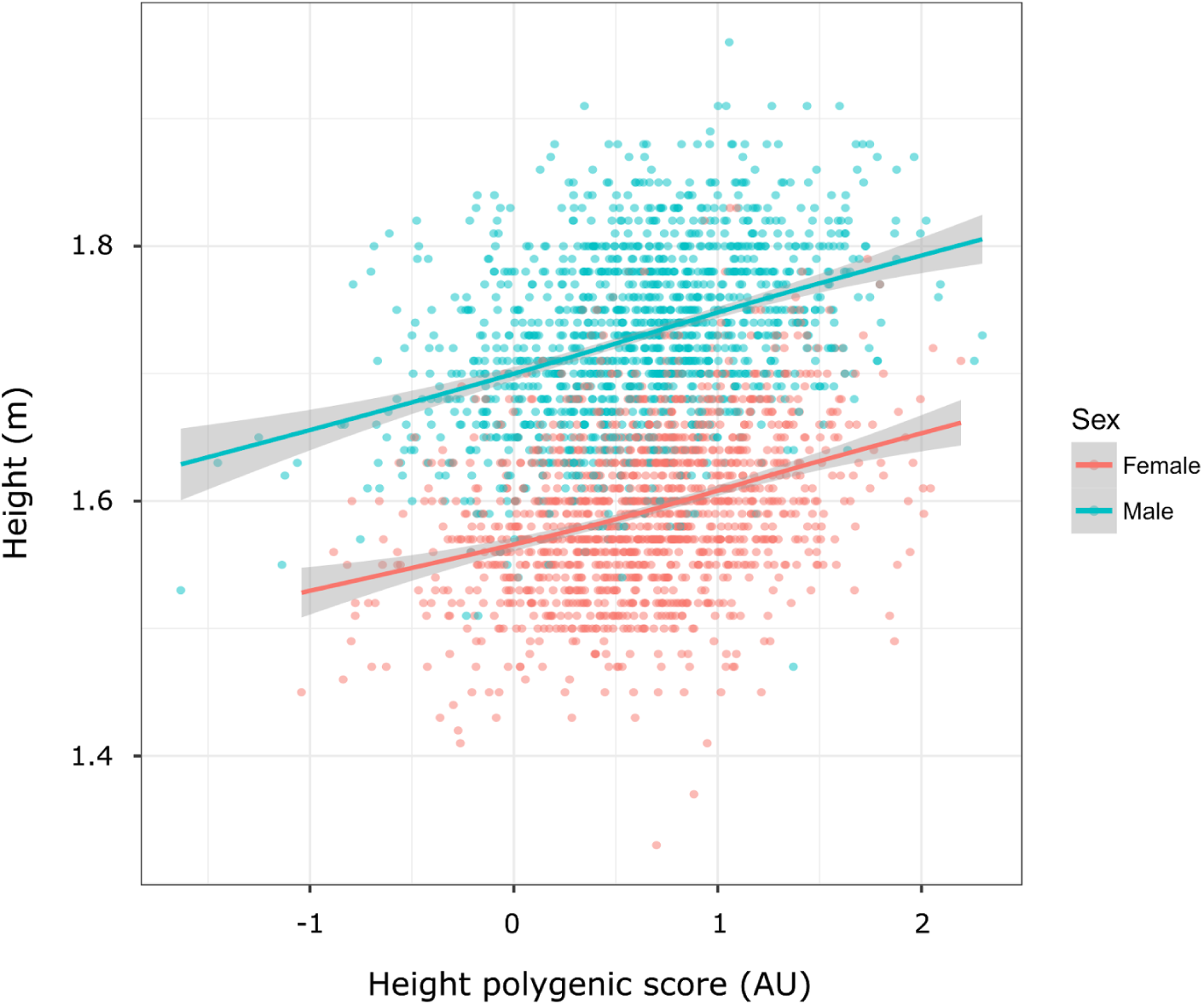
Prediction of height in MGRB using a polygenic score (Wood et al., 2014). Each point represents the predicted and observed height of an MGRB individual; lines denote GCV-penalised generalised additive model thin plate spline fits.

**Supplementary Figure 6:**
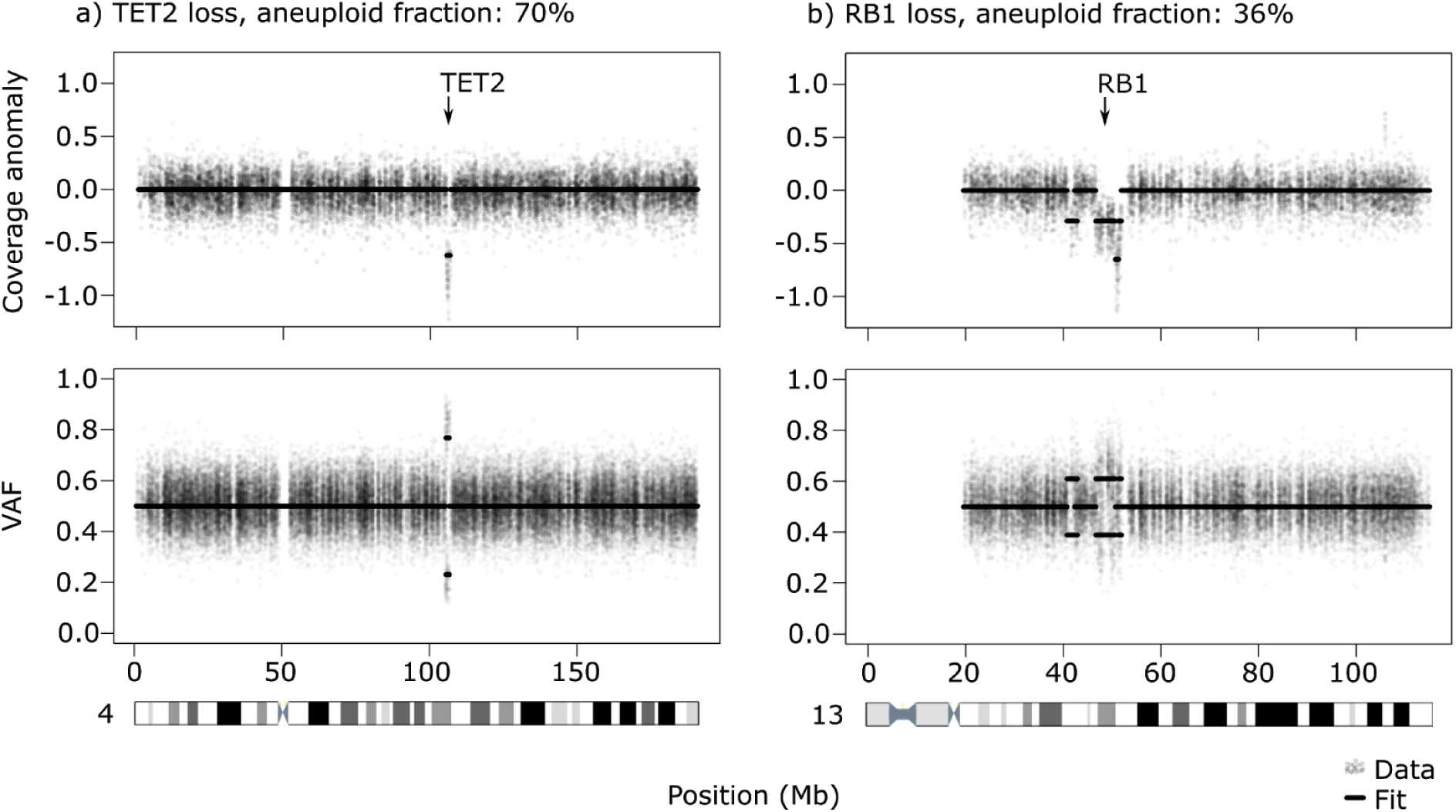
Examples of subclonal copy number variation observed in the MGRB. Figures show background-corrected coverage (top panels) and heterozygous variant allele frequency (bottom panels) as a function of genomic location. Individual locus measurements are represented by semi-transparent dots, with model fits indicated by horizontal segments. These samples demonstrated loss of a single copy of TET2 in an estimated 70% of nucleated blood cells (a), or loss of a single copy of RB1 in approximately 36% of cells (b). Coverage is background-corrected and on a log_2_ scale, with zero indicating diploidy.

**Supplementary Figure 7:**
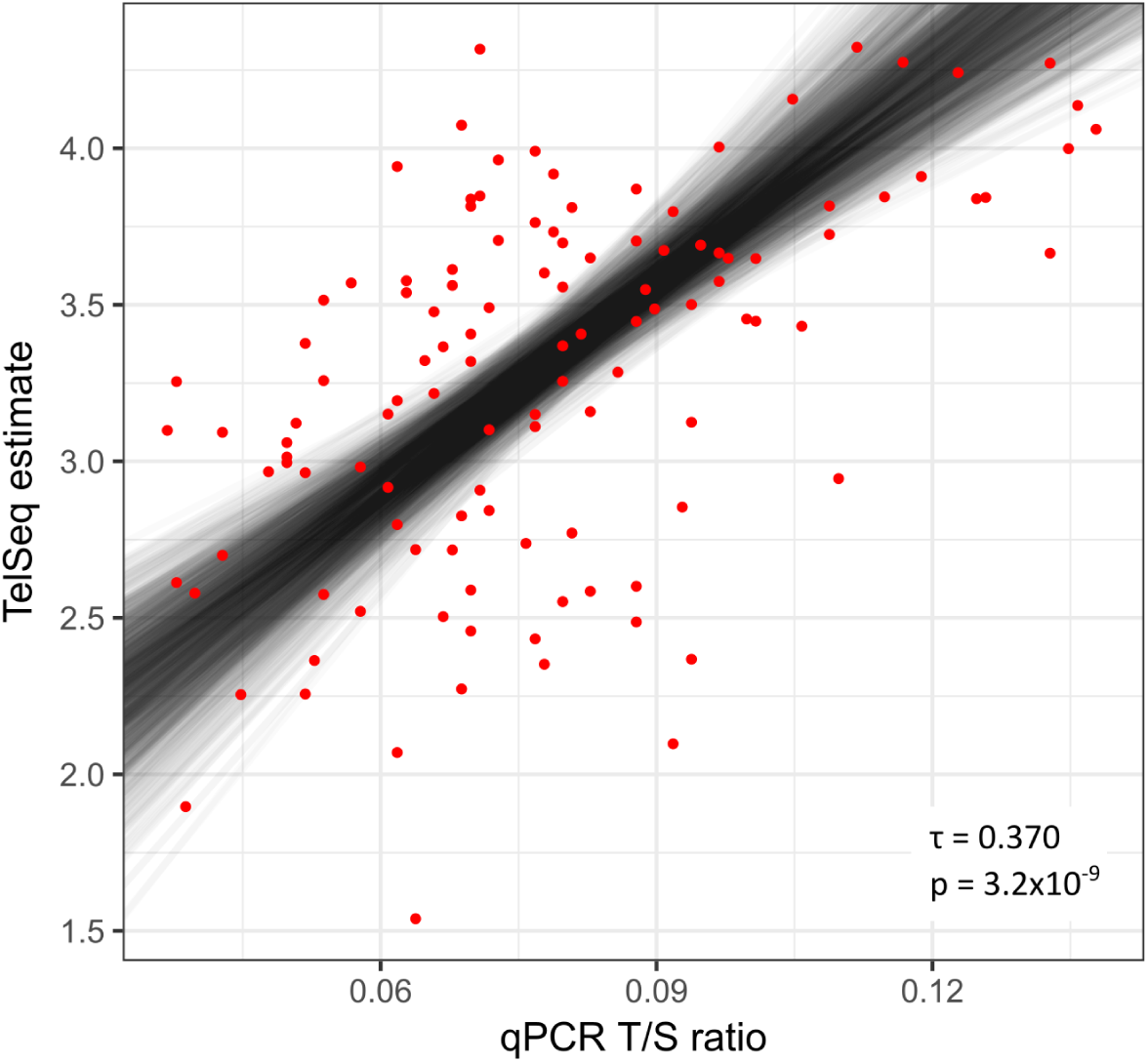
Comparison of Telseq WGS telomere length estimates to qPCR measurements. Points denote 119 randomly-selected samples from the MGRB and ASRB cohorts; one outlier with a Telseq estimate over 5 was excluded. Telseq estimates are directly as reported by the software; qPCR measurements are telomere / single copy gene copy ratios. Lines represent fits from 1000 bootstrap replicates of Deming regression using within-bootstrap median absolute deviation as an empirical variance estimate. The measures are significantly correlated (Kendall’s τ = 0.370, *p* = 3.2 × 10^−9^).

**Supplementary Figure 8:**
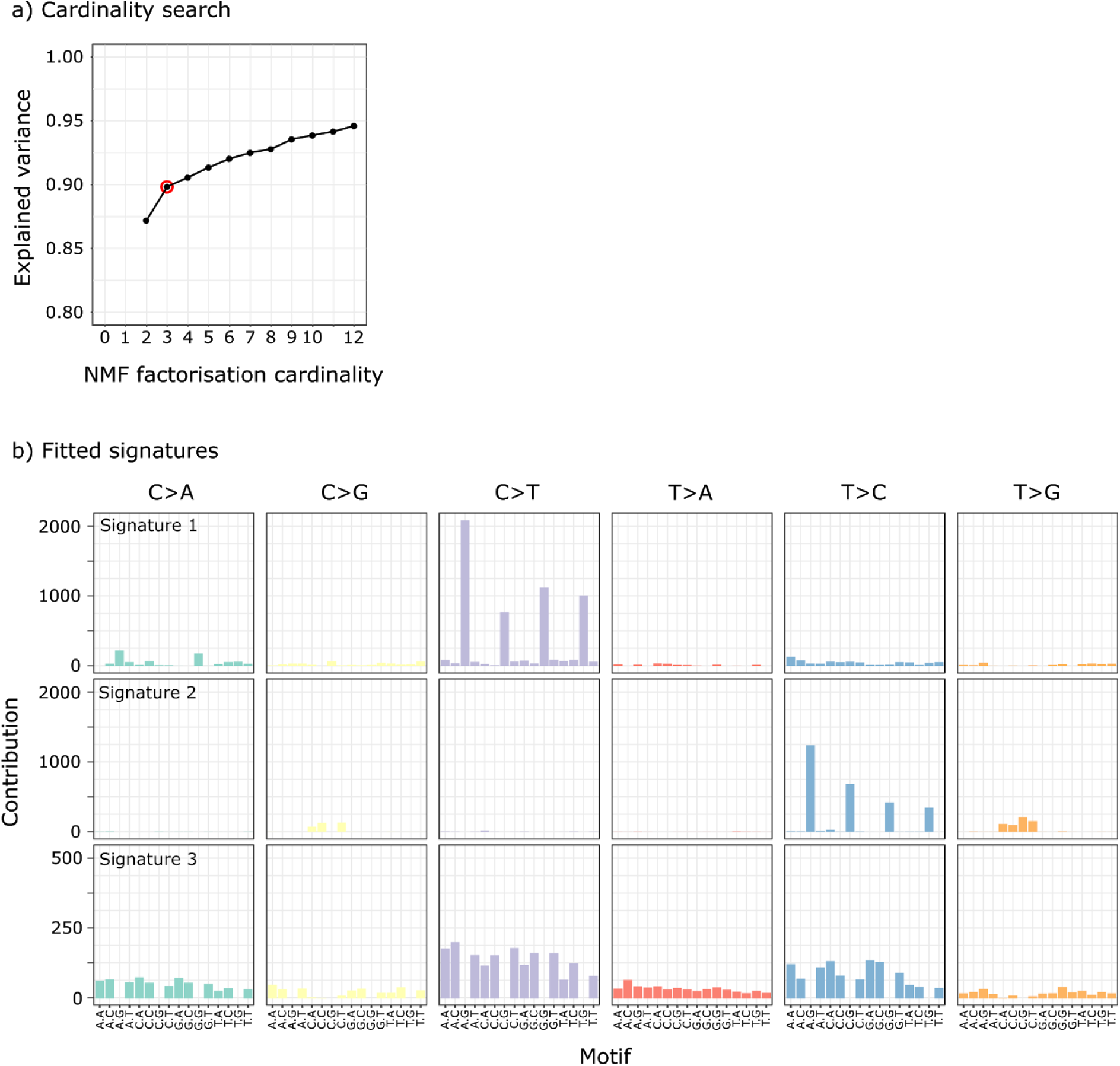
Somatic variant motif factorization. A cardinality search on age-grouped samples indicated that a cardinality of 3 was appropriate, being the inflection point on the explained variance vs cardinality plot (a, selected cardinality marked with red circle). When the single-sample motif frequencies were factorized at this selected cardinality, the three signatures resulting were well resolved (b), with Signatures 1 and 3 respectively resembling Signatures 1 and 5 as previously reported (Alexandrov et al., 2015).

**Supplementary Table 1:** Rates of structural variation (SV) detected in the MGRB. Mean event counts per individual are given, with standard deviation in parentheses.

| *SV class* | *Rate* | *(SD)* | *Fraction* |
| --- | --- | --- | --- |
| <u>*GRIDSS-reported*</u> |  |  |  |
| Insertion | 45 | (7) | 0.5% |
| Deletion | 2750 | (136) | 33.1% |
| Indel | 327 | (21) | 3.9% |
| Duplication | 882 | (98) | 10.6% |
| Inversion | 32 | (5) | 0.4% |
| Total GRIDSS | 4036 | (249) | 48.6% |
| <u>*Mobster-reported*</u> |  |  |  |
| L1 insertion | 1072 | (380) | 12.9% |
| ALU insertion | 2754 | (246) | 33.2% |
| SVA insertion | 436 | (92) | 5.3% |
| HERV insertion | 3 | (2) | < 0.1% |
| Total Mobster | 4264 | (634) | 51.4% |
| Grand total | 8300 | (675) | 100% |

**Supplementary Table 2:** Singleton and polymorphic rates for structural variants (SVs) identified in the MGRB. Structural variant count and fraction are given as a function of the number of samples identified to share that variant. Sample ranges are inclusive.

| *Samples with variant* | *SV count (fraction)* |
| --- | --- |
| 1 (singletons) | 155287 (17.1) |
| 2 - 10 | 589771 (65.0) |
| 11 - 100 | 139340 (15.3) |
| 101 - 500 | 13669 (1.5) |
| 501 - 1000 | 4623 (0.5) |
| 1001 - 1500 | 2178 (0.2) |
| 1501 - 2000 | 1367 (0.2) |
| 2001 - 2500 | 1218 (0.1) |
| 2501 - 2570 | 486 (0.1) |

**Supplementary Table 3:** Structural variants identified in MGRB that may disrupt an ACMG incidentally-reportable gene.

| *Gene* | *Variant* | *Predicted effect* |
| --- | --- | --- |
| *PCSK9* | 1:g.55494888_55509044del | Loss of 5' UTR and exon 1. |
| *SMAD4* | 18:g.48556989_48573287del | Loss of 5' UTR. |
| *TMEM43* | 3:g.141047610_14277101del | Deletion of entire *TMEM43* locus. |
| *VHL* | 3:g.10067411_10421889inv | Inversion of entire *VHL* locus. |

**Supplementary Table 4:** Clinical and demographic characteristics of the 45 and Up cancer cases, compared to the 45 and Up cancer-free individuals included in the MGRB. Cancer cases had some evidence of a cancer diagnosis prior to age 70, either by self report or admission and registry records; cancer-free individuals had no such evidence prior to age 70. Aggregate statistics are medians, with ranges in parentheses. As some individuals had multiple cancers, the sum of cancer types exceeds the number of cancer cases.

| Measure | Cancer cases | Cancer-free |
| --- | --- | --- |
| Individuals | 269 | 717 |
| (percent female) | (45.3%) | (59.3%) |
| Age at collection (years) | 71 | 70 |
|  | (64 – 88) | (64 – 91) |
| Height (m) | 1.70 | 1.66 |
|  | (1.47 – 1.96) | (1.37 – 1.91) |
| Mass (kg) | 76.0 | 72.0 |
|  | (44.5 – 120.0) | (36.0 – 147.0) |
| Cancer type |  |  |
| Prostate | 74 | — |
| Melanoma of skin | 58 | — |
| Colorectal | 40 | — |
| Breast | 26 | — |
| Non-melanoma skin | 20 | — |
| Lung | 13 | — |
| Bladder | 10 | — |
| Other | 124 | — |

